# Time-Resolved Analysis of the Cell Wall Proteome in *Saccharomyces cerevisiae* S288c During Batch Fermentation

**DOI:** 10.64898/2026.01.05.697623

**Authors:** Marie Yammine, Antoine Picavet, Emmanuel Poilpré, Fabrice Bray, Stéphanie Flament, Isabelle Mouly, Christian Rolando

**Author notes:** Correspondance : M.Y. -; C.R. –. Institut Pasteur, Université Paris Cité, and CNRS UAR2024, Mass Spectrometry for Biology, Paris, France.

## Abstract

The yeast cell wall is a highly dynamic and multifunctional structure that is essential for maintaining cellular integrity, protecting against environmental stresses, and enabling adhesion, signaling, and interactions with the surrounding environment. Its chemical composition and organization are strongly influenced by external factors such as temperature, pH, nutrient availability and their delivery mode. In batch culture systems, yeast cells grow in a closed environment with limited nutrients, leading to well-defined growth phases that reflect major metabolic transitions. While global proteomic changes during these phases have been described, the temporal regulation of cell wall protein (CWP) expression remains insufficiently characterized.

In this study, the temporal remodeling of the cell wall proteome of *Saccharomyces cerevisiae* S288c was examined during batch cultivation in rich medium over 24 hours. A classical proteomics workflow was applied to analyze CWPs from samples collected at multiple time points over the cultivation period. The analysis revealed substantial qualitative and quantitative changes in CWPs expression linked to metabolic shifts between growth phases. Proteins involved in cell wall remodeling and glycoprotein biosynthesis were particularly enriched at the initial sampling point (b-T0h), corresponding to the transition from flask cultivation to bioreactor conditions, and overall CWP abundance was highest during this early growth stage.

Time-resolved quantitative, transcription factor, and functional enrichment analyses revealed coordinated regulation of cell wall adaptation. Stationary-phase specific protein markers linked to glucose depletion were identified, offering insight into nutrient-limited remodeling. Comparisons with previous studies showed variability driven by strain differences, culture conditions, and methodological approaches.

## Introduction

*Saccharomyces cerevisiae* has been a valuable microorganism throughout human history, playing a central role in food and beverage fermentation for thousands of years (Sicard & Legras, 2011). Since the complete sequencing of its genome in the late 20th century, and thanks to its well-characterized physiology and long-standing record of safe use, the applications of *S. cerevisiae* have expanded far beyond traditional fermentation (Goffeau et al., 1996; Johnson & Echavarri-Erasun, 2011; Nielsen, 2015). Today, it is also a key player in diverse and advanced biotechnological processes (Botstein & Fink, 2011; Parapouli et al., 2020). In addition to its applied importance, *S. cerevisiae* has profoundly advanced our understanding of eukaryotic biology, serving as a simple, cost-effective, and genetically tractable model organism (Botstein & Fink, 2011; Goffeau et al., 1996; Nielsen, 2019b). Its versatility stems from its well-defined cellular structures and metabolic pathways, which can be exploited in both native and genetically modified forms (Jouhten et al., 2016). Central to all its functions is metabolism, the transformation of nutrients into energy and essential biomolecules for growth and survival (Feist et al., 2009). Consequently, the study of *S. cerevisiae*’s nutritional requirements is fundamental to understanding its physiology, genetics, and industrial potential. Insights into yeast nutrition are crucial for optimizing growth, regulating metabolic activity, improving stress responses, and designing defined media and controlled culture conditions (Broach, 2012). In addition to the consideration of culture medium composition, the culture medium feeding strategy is also crucial for *S. cerevisiae* yeast growth. There are four major culture medium feeding strategies: discontinuous or “batch”, semi-continuous or “fed-batch”, continuous and perfusion (Lidén, 2002; Paulová et al., 2013).

In batch culture, all nutrients required for cell growth are initially provided in the medium, and the culture is initiated by inoculating biomass from a seed reactor. The system is closed, with no addition or renewal of nutrients during the process (Rajpurohit & Eiteman, 2022). Cell growth varies over time, driven by nutrient availability and consumption and concomitant metabolic shifts (Nielsen, 2019a). After an initial lag phase (during which cells adapt to the new environment), growth enters an exponential (log) phase, which slows down at the diauxic shift as preferred nutrients (e.g., glucose) become depleted (Alberghina et al., 2012). This is followed by the stationary phase, where growth ceases and a balance between cell division and death is reached (Werner-Washburne et al., 1993). Eventually, the culture enters the death phase, driven by nutrient exhaustion and accumulation of waste products (Gonzalez & Aranda, 2023). Once nutrients are fully consumed, the process halts, inherently limiting both productivity and scalability for industrial applications (Gonzalez & Aranda, 2023; Paulová et al., 2013; Rajpurohit & Eiteman, 2022). These dynamic physiological transitions are closely linked to changes at multiple omics levels (Zampar et al., 2013), including the transcriptome (Gombert et al., 2001; Kolkman et al., 2005), proteome (den Ridder et al., 2023; Kolkman et al., 2005; Murphy et al., 2015), and metabolome (Gombert et al., 2001; Tu et al., 2007), each reflecting the distinct phases of growth under defined culture conditions. Recent advances have leveraged machine learning and high-dimensional proteomics data to construct dynamic models that capture yeast behavior over time across various culture environments (Moimenta et al., 2025; Olivares□Hernández et al., 2010). These integrated approaches not only account for uncertainty but also identify key proteins most relevant to growth dynamics.

Despite numerous studies investigating proteome dynamics during yeast culture under various environmental conditions and setups (Costenoble et al., 2011; de Groot et al., 2007; Picotti et al., 2009), including organelle-specific analyses (Di Bartolomeo et al., 2020; Helbig et al., 2009; Sickmann et al., 2003), the temporal evolution of the cell wall proteome (CWP) of *S. cerevisiae* during batch culture has not, to our knowledge, been systematically described. Previous work has demonstrated that the biochemical composition of the yeast cell wall is subject to culture-dependent remodeling, underscoring its dynamic nature (Aguilar□Uscanga & Francois, 2003). Our interest in this organelle stems from its critical role as a rigid barrier that both protects the cell and mediates interactions with the external environment (Hapala et al., 2013; Osumi, 1998). Moreover, the yeast cell wall is composed of diverse biomolecules (Orlean, 2012), such as mannoproteins, β-glucans, and chitin, that are functionally and structurally valuable for a wide range of applications (Jofre et al., 2024).

In this study, we aimed to characterize the temporal remodeling of the yeast cell wall proteome during the progression of a batch culture in a bioreactor using a rich medium. The model organism used was *Saccharomyces cerevisiae* strain S288c, cultivated in rich medium. Cell samples were collected at key growth phases - exponential (log) phase, diauxic shift, and stationary phase - for yeast cell wall (YCW) extraction. The extracted fractions were then analyzed using label-free bottom-up quantitative proteomics to profile dynamic changes in cell wall protein composition over time.

## Experimental procedures

### Yeast culture conditions

*Saccharomyces cerevisiae* S288c strain (ATCC: 204508, MATα SUC2 gal2 mal2 mel flo1 flo8-1 hap1 ho bio1 bio6) seed was grown for 24 hours at 30 °C in an Erlenmeyer flask (250 mL) with sterilized YPD medium (150 mL, 1% Gibco^TM^ Yeast Extract, 2% Gibco^TM^ bactopeptone, and 2% glucose from Carlo Erba reagents) under agitation at 120 rpm.

A yeast suspension at 5 g dry weight/L was obtained after centrifugation of the seed culture and washing of the pellet, from which 500 µl were then transferred into a 6 L round-bottom flask filled with 3 L of sterilized standard YPD medium. Following 16 hours incubation (corresponding to the exponential phase) at 30 °C with shaking at 120 rpm, the seed was centrifuged and washed once with physiological water to generate a cream concentrated at 8 g/L. A 7 L bioreactor (Applikon Biotechnology ez-Control autoclavable Bioreactor 7 L) containing 5.3 L of sterilized YPD medium was inoculated with 0.45 g (yeast dry matter) of the cream. The batch bioreaction was carried out in three independent experiments, for 24 hours at 30 °C with an airflow of 1 VVM (volume of air per unit volume of growth medium per minute), and the medium pH was controlled at 5.0 by the automatic addition of 10% H_2_SO_4_ and 10% NaOH. During the bioreaction, samples were taken to realize spectrophotometric OD measurements of cell growth at 600 nm using a UVILINE 8100 SECOMAM spectrophotometer. The spent medium containing the cells was filtered and served to determine the concentrations of glucose and ethanol by HPLC-H+ ion exchange chromatography method using Prominence HPLC system (Shimadzu) equipped with Aminex HPX-87H (Catalog number: 1250140, Pkg of 1, 300 x 7.8 mm;Biorad, CA, USA). In addition to the seed (T = 0 h), four samples each of 1.25 g dry matter, corresponding to different studied growth phases, were harvested during the culture: after 4 h (mid-exponential phase), 7.5 h (early diauxic phase), 9 h (late-diauxic shift) and 24 h (stationary phase).

### YCW isolation by mechanical disruption

Mechanical disruption was performed as described previously (Yammine et al., 2022). Yeast cells were resuspended at 400 mg/mL in ice-cold lysis buffer (10 mM Tris-HCl pH 7.5, 1X cOmplete^TM^ protease inhibitor cocktail: #11697498001, Roche Diagnostics, Germany), and transferred to BeadBug™ triple-pure tubes prefilled with glass beads (0.5 mm diameter, Benchmark Scientific, NJ, USA). 15 cycles of 1 min – homogenization and 5 min – resting on ice were applied using BeadBug™. Subsequently, lysates were separated from beads, and three washing steps were carried out for beads with a solution of 1 M NaCl. Lysates and washing solutions were combined and centrifuged for 10 min (4000 g, 4 °C, Eppendorf™ Centrifuge 5430 R). To decrease the cytosolic proteins contaminants, three successive washings of the pellet were performed with 1 M NaCl solution. To dissolve the membranous organelles, the pellet was extracted twice with boiling SDS buffer (2% SDS, 100 mM β-mercaptoethanol, 100 mM EDTA, 150 mM NaCl in 50 mM Tris-HCl pH 7.5) for 10 min each, followed by a 5 min-centrifugation step at 20 000 g (Beckman Coulter Allegra™ 64R Centrifuge). The extracted pellet was further washed for multiple times with 1.5 mL ultrapure water to result in YCW isolates, that were dried under vacuum in an Eppendorf™ Concentrator plus (Concentrator Savant ISS110).

### Proteomics sample preparation and LC-MS/MS analysis

Protein concentration in YCW isolates was determined using Pierce™ Bicinchoninic acid protein assay kit (Thermo Scientific^TM^). 50 µg of proteins from each sample were digested overnight at 37 °C with trypsin (1 µg, V5111, Promega) according to the bottom-up proteomics workflow using eFASP method (Erde et al., 2014) in filtration devices (Amicone® 10 kDa MWCO, Millipore). Peptides were recovered and extracted three times with ethyl acetate (270989; Sigma-Aldrich, Saint-Louis, MO, USA) and dried in Eppendorf™ Concentrator plus. Peptides concentration was determined after adding 10 μL of 0.1% formic acid using the absorbance measurement at 214 nm with a spectrophotometer (Denovix® DS-11 + spectrophotometer; Denovix Inc., USA).

A nanoflow HPLC instrument (U3000 RSLC Thermo Fisher Scientific^TM^) was used, coupled on-line to a Q Exactive plus mass spectrometer (Thermo Scientific^TM^) with a nanoelectrospray ion source. Using partial loop injection, the preconcentration trap (Thermo Scientific^TM^, Acclaim PepMap100 C18, 5 µm, 300 µm i.d × 5 mm) was loaded for 5 min at 10 µL.min^−1^ with1 µg of peptides and buffer A (5% acetonitrile and 0.1% formic acid). Separation was carried out with a reversed phase column (Thermo Scientific^TM^, Acclaim PepMap100 C18, 3 µm, 75 mm i.d. × 500 mm) heated at 45 °C at a flow rate of 250 nL.min^−1^ using a 160 min linear gradient of 5–50% buffer B (75% acetonitrile and 0.1% formic acid). The column washing step was performed for 10 min with 99% of buffer B followed by a reconditioning with buffer A. The total time for an LC-MS/MS run was about 180 min long.

### Proteomics data analysis

The acquired raw files were analyzed with Proteome Discoverer^TM^ 2.2 software (Thermo Scientific^TM^) with Sequest search engine against *Saccharomyces cerevisiae* S288C strain dataset (orf_trans_all) from Saccharomyces Genome Database (SGD) (modified in January 2015). Mass tolerance of peptide was specified at 10 ppm for MS and 0.01 Da for MS/MS. Fixed modifications as cysteine carbamidomethylation and variable modifications as methionine oxidation were included. Proteins were identified with at least two unique peptides. A label free quantification method was implemented in data processing using Minora algorithm. Gene Ontology (GO) analysis using Uniprot Knowledgebase (UniprotKB) and SGD GO Slim mapper tool was performed, specifically in analyses related to cellular component category permitting to study proteins’ subcellular location. An in-house database for yeast cell wall proteins (CWP) was also implemented. Additionally, the data were compared with proteomic profiles derived from a fed-batch culture conducted with the same strain and culture conditions, as previously reported (Yammine et al., 2022). YEASTRACT+ (Monteiro et al., 2019; Teixeira et al., 2022) analysis was performed using the “Rank by TF” function, evaluating all activating and repressing transcription factors. Venn diagrams were traced using jvenn tool (Bardou et al., 2014) and venny 2.1.0 (https://bioinfogp.cnb.csic.es/tools/venny/). Heatmaps for CWPs expression profiles, hierarchical clustering by Euclidean distance and the correlation between CWPs were established using Morpheus (https://software.broadinstitute.org/morpheus/), based on z-score normalization (to average and standard deviation) and Spearman rank correlation function respectively.

### Experimental design and statistical rationale

For our differential proteomics investigation during a batch culture kinetics of *Saccharomyces cerevisiae* in rich medium, we performed three independent biological replicates from three separate inoculated seeds. 5 samples were withdrawn at the same time points from the three independent seeds and bioreactors. Sample processing was performed in parallel under identical conditions for technical variability minimization. No technical replicates were performed for proteomics as previous studies have shown that the LC-MS analyses produced only negligible variations (Tabb et al., 2010). Identified proteins were analyzed using Perseus software (v.2.1.5.0) and a protein was considered present in at least 2 out of replicates. Protein abundances were normalized. Statistical significance was determined using Student’s *t*-test and ANOVA test with Benjamini–Hochberg correction. Overexpression is considered if the log2 fold change between two conditions was greater than 1.5 and the statistical significance was defined as p < 0.05.

## Results

### Identification and Quantification of Proteins in Batch Bioreactor Culture

*S. cerevisiae* S288c reference strain was specifically chosen for its well-characterized genome and the comprehensive omics data available, facilitating thus data integration and interpretation. Batch cultures were sampled across all growth phases, spanning from proliferation to stationary-phase arrest (supplemental Fig. 1). Sampling time points correspond to mid-exponential phase (b-T4h), early diauxic phase (b-T7.5h), late-diauxic phase (b-T9h) and stationary phase (b-T24h), in addition to a sample from the seed that corresponds to the beginning of bioreactor culture (b-T0h). To ensure the high data representativeness and robustness and to capture biologically relevant changes, independent batch fermentations were conducted in triplicate bioreactors under the strict controlled environmental conditions, including oxygen availability, pH and temperature. Indeed, this robustness is evidenced by a Pearson correlation coefficient *r* greater than 0.8 between the replicates across all conditions (supplemental Table S1). Under aerobic conditions, it is well known and established that glucose-grown cultures of *S. cerevisiae* engage in both respiration and fermentation, resulting in the accumulation of ethanol and other fermentation products. This is followed by a diauxic growth phase, where these metabolites are subsequently respired until stationary phase is reached. This is clearly shown in the growth curve (supplemental Fig. 1A) as well as glucose consumption and ethanol production profiles in our study (supplemental Fig. 1B).

Following YCW extraction and trypsin digestion, peptide samples were analyzed by bottom-up label free quantification proteomics approach using a 3 h LC gradient. Over the course of batch fermentation, the number of quantified proteins slightly increased from an average of 950 at b-T0h to 1038 at b-T24h (Fig. 1A). A protein was considered quantified at a given growth phase if it was detected in at least two replicates under the same condition. The sets of quantified proteins were compared across all growth phases (Fig. 1B) using jvenn tool (Bardou et al., 2014). When combined, all datasets yield a matrix comprising a total of 1,558 unique quantified proteins. As illustrated in the Venn diagram (Fig. 1B), 410 proteins were common to all phases, while 544 proteins were unique to specific phases, primarily at b-T0h (274 proteins) and b-T24h (183 proteins). These findings indicate that the proteomes at the start and end of fermentation are more distinct from those observed during the active growth phase. This observation is further supported by the correlation matrix presented in supplemental Table S1. The proteomic profile of the late diauxic phase (b-T9h) shows moderate correlation with both the mid-exponential phase (b-T4h) and the early diauxic shift (b-T7.5h), with correlation coefficients ranging from 0.6 to 0.7. Notably, b-T0h exhibits the weakest correlation with all other phases (correlation coefficient < 0.6). In contrast, the early diauxic shift phase (b-T7.5h) correlates strongly with both the mid-exponential (b-T4h) and late diauxic (b-T9h) phases, with coefficients exceeding 0.8. Additionally, the late diauxic phase (b-T9h) also shows a good correlation (coefficients between 0.75 and 0.8) with both the mid-exponential (b-T4h) and stationary (b-T24h) phases. To gain insight into their cellular localization, we analyzed the Gene Ontology (GO) annotations of the top 100 most abundant proteins quantified in each replicate and condition. These proteins account for over 80% of the total protein abundance (data not shown) and their distribution (in terms of relative abundance) is presented in Figure 1C. We observed an increase in the abundance of mitochondrial proteins and cell wall proteins (CWPs), alongside a decrease in ribosomal proteins, over the course of batch culture growth. This trend highlights the metabolic shift from a respiro-fermentative growth phase, characterized by high protein synthesis activity, to a respiratory state during the diauxic phase, where mitochondrial activity is predominant.

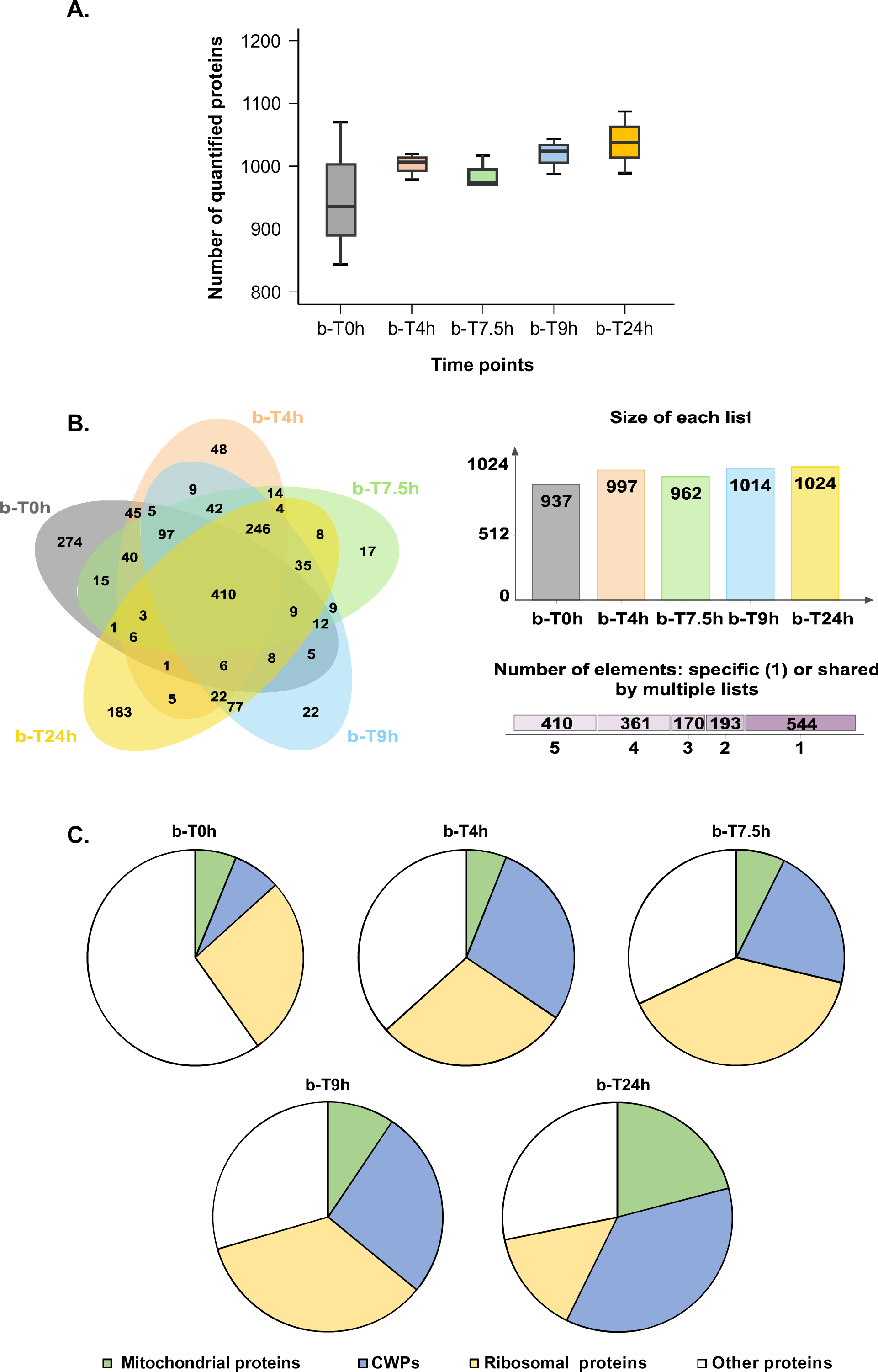

Notably, the CWPs among the top 100 most abundant proteins include several moonlighting proteins, such as Tdh3p, Tdh2p, Tdh1p, Ssa1p, and Ssa2p. These proteins are known to localize to multiple cellular compartments and play roles not only in cell wall structure, but also in glycolysis and stress response. Therefore, we conducted a more in-depth functional analysis of our dataset to better understand the overall proteome dynamics - and specifically the behavior of CWPs - throughout batch fermentation in bioreactors, as detailed subsequently.

### Proteomic Changes Driven by Nutrient Availability in Batch Bioreactor Culture

To comprehensively capture protein expression dynamics throughout the bioreactor batch culture, we conducted both pairwise and multivariate comparisons of samples collected at different time points.

Multivariate analysis of the dataset revealed clear clustering of biological replicates for each time point, highlighting the high reproducibility of the data (Fig. 2A). Principal component analysis (PCA) performed using Perseus software further showed that the inoculum (b-T0h) and the stationary phase (b-T24h) were distinctly separated from the mid-exponential (b-T4h) and early diauxic (b-T7.5h) phases. The late diauxic phase (b-T9h) clustered between the actively growing mid-exponential phase and the stationary phase, reflecting its transitional nature (Fig. 2A).

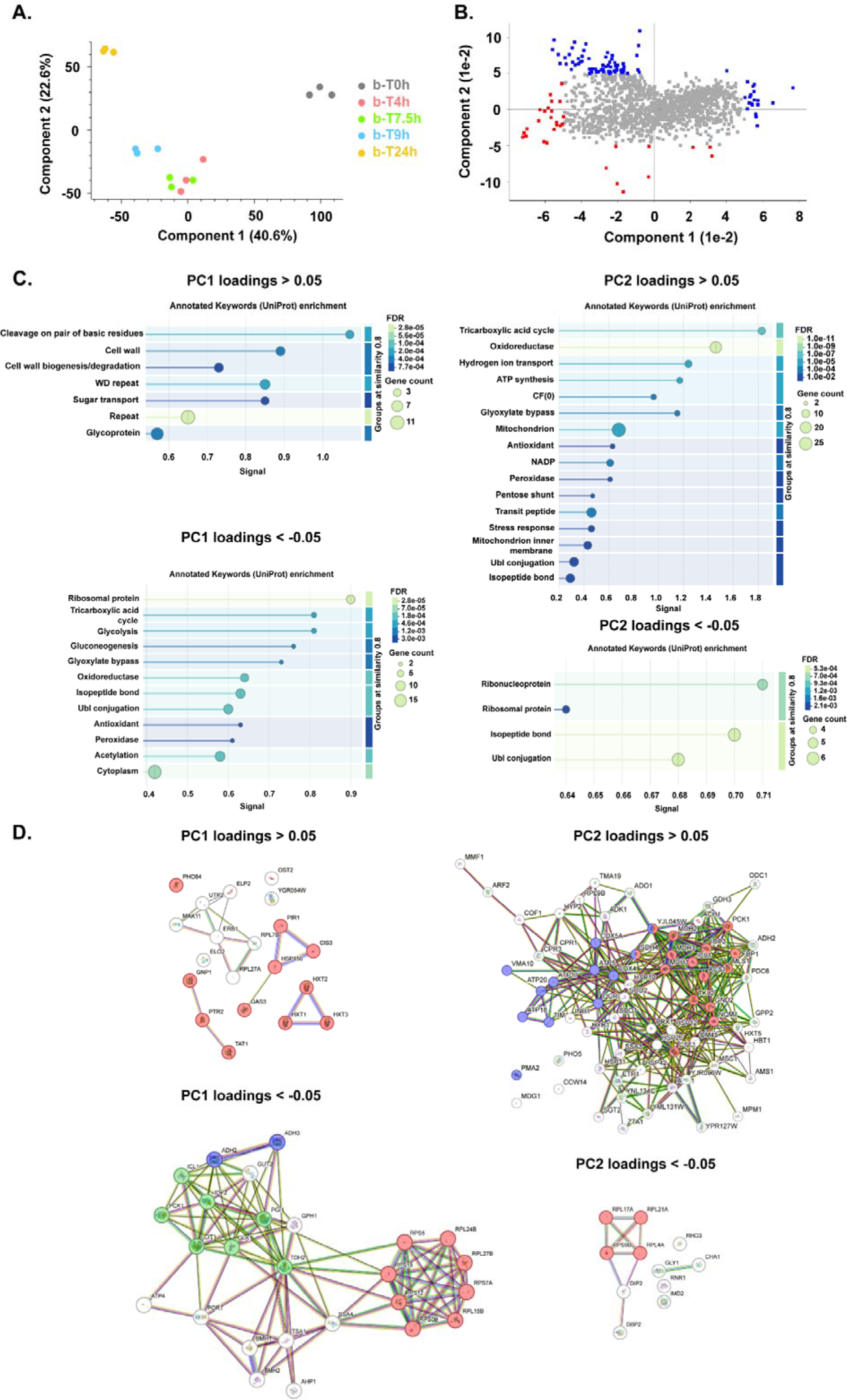

Principal component 1 (PC1), which accounts for 40.6% of the total variance, primarily separated the flask-grown inoculum (b-T0h) from the bioreactor samples (b-T4h to b-T24h). Principal component 2 (PC2), explaining 22.6% of the variance, distinguished the stationary phases, characterized by growth on non-fermentable carbon sources, from the active growth phases on glucose, including the mid-exponential and early diauxic stages (Fig. 2A). We analyzed the differentiating loads (|score| > 0.05) retrieved from Perseus software, highlighted in blue and red in Fig. 2B. The functional analysis of these loads using String DB (Annotated Keywords Uniprot) showed the enrichment of terms related to cell wall and glycoprotein for loads of component 1 whose score > 0.05 and are more enriched in flask batch culture, while oxidoreductase, tricarboxylic acid cycle, gluconeogenesis and ribosomal proteins are enriched for loads of component 1 whose score < - 0.05 that pull towards the bioreactor batch culture (Fig. 2C and 2D). For the loads of component 2, those with as score > 0.05 were enriched in mitochondria, stress response and oxidoreductase, and are more enriched in stationary phase time points, while those with as score < -0.05 were enriched in ribonucleotides and ribosomal proteins in the early growth phases (Fig. 2C and 2D).

For pairwise comparisons, differentially expressed proteins between time points were identified using Student’s *t*-tests and visualized via Volcano plots. The results are summarized in Table 1. The inoculum sample (b-T0h) exhibited a substantial number of overexpressed proteins compared to later time points, ranging from 452 (relative to the mid-exponential phase, b-T4h) to 628 (relative to the stationary phase, b-T24h) proteins. Functional analysis of the 571 proteins consistently overexpressed in b-T0h, using String-DB, revealed enrichment in 12 KEGG (Kyoto Encyclopedia of Genes and Genomes) pathways, including those involved in lipid, *N*-glycan, and *O*-glycan biosynthesis, processes associated with cell wall integrity (supplemental Fig. S2A).

**Table 1.** Number of differentially expressed proteins for pairwise comparison of proteome according to the growth phase in batch culture.

|  | b-T0h | b-T4h | b-T7.5h | b-T9h | b-T24h |
| --- | --- | --- | --- | --- | --- |
| b-T0h |  |  |  |  |  |
| b-T4h | 452<br>382 |  |  |  |  |
| b-T7.5h | 554<br>381 | 0<br>0 |  |  |  |
| b-T9h | 573<br>454 | 46<br>78 | 3<br>30 |  |  |
| b-T24h | 628<br>587 | 342<br>352 | 379<br>404 | 293<br>249 |  |

The proteins were considered significantly differential for a false discovery rate $FDR < 0.05$ and a $S0$ value of 0.1, using Student's $t$ -test. Each cell of the matrix is subdivided into two, figuring two numbers with different cell colors, and indicate a comparison between two time points: one on the matrix row and the other on the matrix column. The numbers written in black in white cells denote the proteins that are overexpressed in time points on the matrix row, whereas those written in black in grey cells correspond to the proteins overexpressed in the time points on the matrix column for each comparison. Black cells correspond to the comparison between the same time points. The cells were filled on one side of the matrix for the sake of simplicity and clarity.

Conversely, the number of proteins downregulated at later time points compared to b-T0h ranged from 381 (relative to the early diauxic phase, b-T7.5h) to 587 (relative to the stationary phase, b-T24h). Functional analysis of the commonly downregulated proteins indicated enrichment in 49 KEGG pathways, primarily related to metabolic processes (supplemental Fig. S3A).

No significant differences in protein expression were observed between the mid-exponential phase (b-T4h) and the early diauxic shift (b-T7.5h), in contrast to the late diauxic shift (b-T9h), which exhibited a greater number of differentially expressed proteins (Table 1). This likely reflects a temporal lag between transcriptional reprogramming and its translational impact during the diauxic shift, a phase characterized by molecular transitions driven by nutrient depletion between the mid-exponential and stationary phases. Functional enrichment analysis of commonly overexpressed proteins revealed 11 and 17 enriched KEGG pathways for the mid-exponential and early diauxic phases respectively (supplemental Fig. S2B and 2C), including pathways related to ribosomes and amino acid biosynthesis. Meanwhile, the analysis of commonly downregulated proteins indicated enrichment in 14 and 20 KEGG pathways (supplemental Fig. S3B and C), involving metabolism, oxidative phosphorylation, and peroxisome functions. These findings suggest a period of high translational activity coupled with reduced respiratory activity at this stage of the culture.

Consistent with previous findings, only minimal changes were detected between the early and late diauxic phases, with just 3 proteins overexpressed in b-T7.5h and 30 in b-T9h (Table 1). The functional analysis demonstrated the enrichment of 28 KEGG pathways for the commonly overexpressed proteins (supplemental Fig. S2D) and the 3 KEGG pathways for the commonly downregulated proteins in the late diauxic shift (b-T9h) (supplemental Fig. 3D), all related to metabolic reprogramming.

As presented in Table 1, the stationary phase (b-T24h) showed a significant number of differentially expressed proteins compared to all other time points. The smallest differences were observed in comparison to the late diauxic shift (b-T9h), with 249 proteins upregulated and 293 downregulated in b-T24h. In contrast, the largest differences were seen relative to the initial time point (b-T0h), with 587 proteins upregulated and 628 downregulated in b-T24h. Functional enrichment analysis of proteins commonly overexpressed in b-T24h revealed 38 KEGG pathways, primarily associated with metabolic processes such as oxidative phosphorylation and gluconeogenesis (supplemental Fig. S2E). Meanwhile, the commonly downregulated proteins in b-T24h were enriched in 16 KEGG pathways, including those related to ribosomal function and RNA metabolism and cell wall integrity (supplemental Fig. S3E).

Then, we compared the upregulated proteins for a specific time point compared to others (supplemental Table S2, S2.11 - S2.15), as well as the downregulated proteins (supplemental Table S2, S2.16 - S2.20). The lists of common proteins were analyzed using YEASTRACT+ (Monteiro et al., 2019; Teixeira et al., 2022) to find transcription factors that regulate their expression (supplemental Table S2, S2.21 - S2.25 and S2.26 – S2.30). Following this analysis, the transcription factor lists were compared for all the upregulated proteins (195 – 210 transcription factors in supplemental Table S2, S2.31) in different time points, as well as for those that were downregulated (200 – 207 transcription factors in supplemental Table 2, S2.32). We were very interested to specific transcription factors, that were not shared between all the time points. The results showed that most transcription factors (181) were common between upregulated and downregulated proteins for different time points, and that only 6 transcription factors were specific to 91 proteins downregulation, and 13 were specifically upregulating 154 proteins (in supplemental Table S2, S2.33 and S2.34). Functional enrichment analysis showed that most of the upregulated proteins were located on the cell periphery and the mitochondria (supplemental Table S2, S2.34 - supplemental Fig. S4A) and those of the downregulated proteins were in the nucleosome (supplemental Table S2, S2.34 - supplemental Fig. S4B). The comparison showed that 35 proteins were up- and down- regulated by specific transcription factors, and were enriched in 10 KEGG pathways, mostly related to metabolism (supplemental Table S2, S2.34 - supplemental Fig. S4C). Among these specific transcription factors, two of them still are not characterized (YKL222C and YPR196W) and are responsible of upregulating 30 proteins

### Time-Dependent Dynamic Expression of Cell Wall Proteins in Batch Bioreactor Culture

The functional enrichment of terms related to cell wall, such as cell periphery, glycoproteins and glycans biosynthesis led us to examine in great details the temporal expression profile of these proteins, in the form of heatmaps. The heatmaps were constructed using Morpheus (https://software.broadinstitute.org/morpheus/), based on z-score normalization (to average and standard deviation) (data from supplemental Table S3).

Hierarchical clustering using Euclidean distance was performed based on CWP abundances. In total, 44 CWPs were quantified in our dataset, including 17 proteins that are also known to localize to other organelles (shown in blue in Supplementary Table S3 and in regular font in Fig. 3). As illustrated in Fig. 3A, replicates from each sampling time point clustered tightly together, except for one replicate each for b-T7.5h and b-T9h, both corresponding to the metabolic transition phase of the culture. The b-T0h samples formed a distinct cluster that was clearly separated from all other time points (Fig. 3).

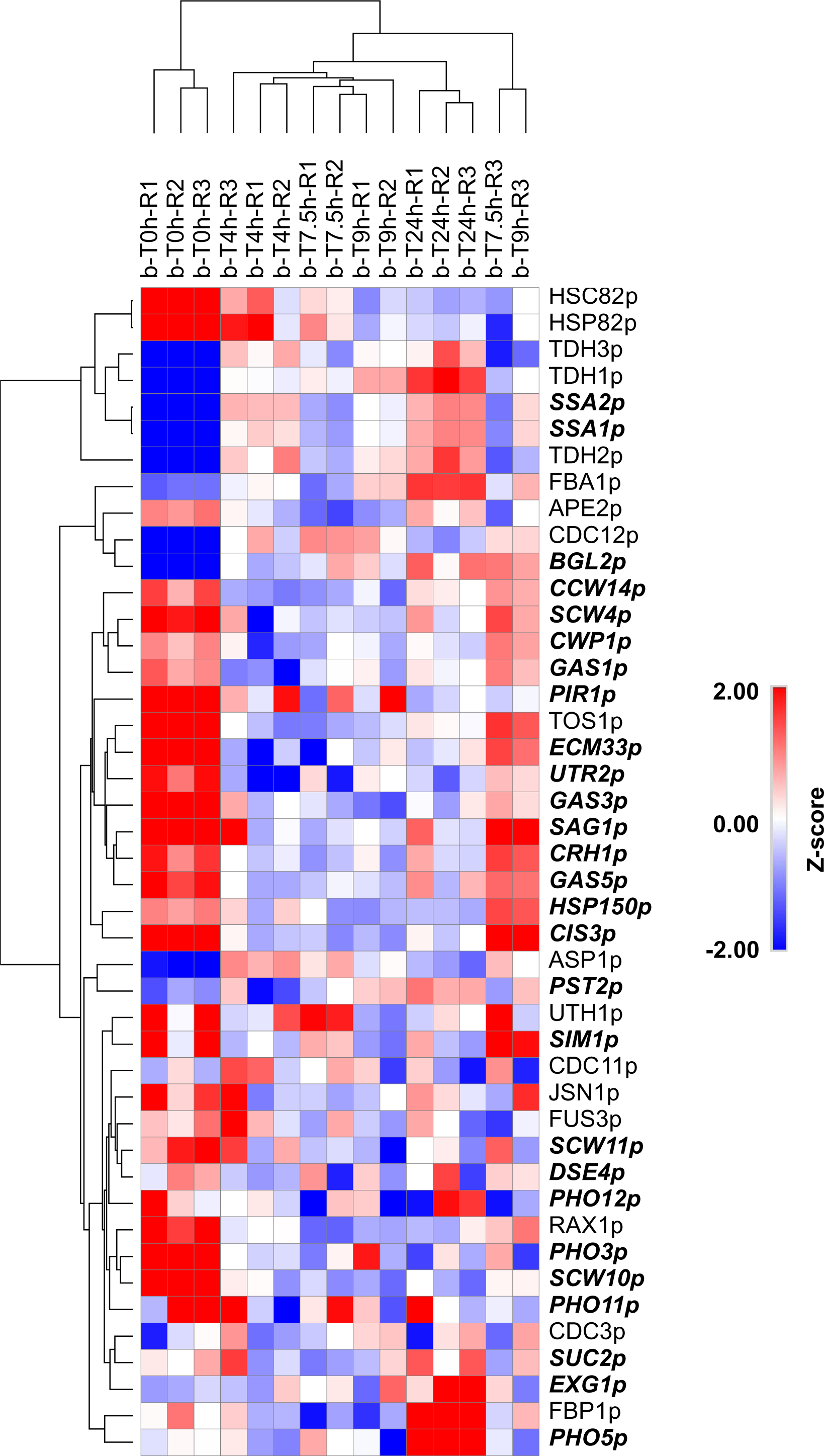

At b-T0h, 27 CWPs were highly expressed (in red), while 10 were downregulated (in blue) compared to the subsequent time points. Notably, the expression of these 10 downregulated proteins increased as the batch cultivation progressed in the bioreactor. Most of these proteins (Tdh3p, Tdh1p, Tdh2p, Cdc12p, Asp1p, Cdc3p, Cdc11p, Ssa1p, and Ssa2p) are associated with multiple organelles, except for Bgl2p, which is exclusively localized to the yeast cell wall (Fig. 3).

Row clustering revealed two major protein groups. The first cluster contains seven proteins (Hsc82p, Hsp82p, Tdh1p, Tdh2p and Tdh3p), all of which may localize outside the cell wall. Within this cluster, Hsp82p and Hsc82p displayed high expression levels at b-T0h that steadily decreased throughout the culture, reaching their lowest abundance at the end of fermentation. In contrast, the remaining proteins in this cluster showed the opposite trend: minimal expression at b-T0h followed by a progressive increase, peaking at b-T24h. The second cluster is subdivided into 3 subclusters: the first subcluster included 4 proteins (Fba1p, Ape2p, Cdc12p and Bgl2p), whose expression varied across the batch culture and mainly increased during the transition phase (b-T7.5h and b-T9h). Fba1p and Bgl2p expression peaked at b-T24h unlike Cdc12p, whereas Ape2p expression peaked at b-T0h. The second subcluster contained 14 proteins (***Ccw14p, Scw4p, Cwp1p, Gas1p, Pir1p,*** Tos1p, ***Ecm33p, Utr2p, Gas3p, Sag1p, Crh1p, Gas4p, Hsp150p, and Cis3p***), 13 of which are exclusively localized to the yeast cell wall (shown in bold and italic in Fig. 3). These proteins were strongly upregulated at b-T0h and are primarily involved in yeast cell wall structure and remodeling. The remaining 19 CWPs exhibited more complex expression profiles. Of these, 10 were upregulated in b-T0h (Uth1p, ***Sim1p***, Jsn1p, Fus3p, ***Scw11p, Dse4p***, Rax1p, ***Pho3p, Scw10p, and Pho11p***), with five exclusively localized to the YCW (highlighted in bold and italics in Fig. 3). 6 proteins were specifically upregulated at the stationary phase (b-T24h) during the batch culture in bioreactor (Fig. 3). These proteins are Fba1p, Fbp1p, Pst2p, Suc2p, Pho5p and Exg1p, mostly implicated in carbon metabolism when fermentable carbon sources are limited. During the transition diauxic phase, the expression of some proteins is increased compared to the other time points of the batch culture, including ***Uth1p, Hsp150p, Cwp1p, Gas1p and Sim1p*** (Fig. 3). During the exponential growth, the majority of CWPs were downregulated (Fig. 3). Spearman rank correlation of CWPs expression showed 29 CWPs whose expression was positively correlated (correlation coefficient > 0.5, red bars in supplemental Fig. S5), mostly containing proteins that are strictly located to the YCW, whereas 12 CWPs were negatively correlated (correlation coefficient < -0.5, blue bars in supplemental Fig. S5), mostly containing proteins that can be in other organelles.

To provide a clearer visual overview of each CWP, we generated kinetic plots by averaging normalized abundances for each CWP of the biological replicates at each time point and expressing them relatively to the total abundance of the CWP across the time points. The corresponding standard deviations were also displayed. We also differentiated CWPs found exclusively in the YCW (Fig. 4) from those present in additional organelles (Fig. 5).

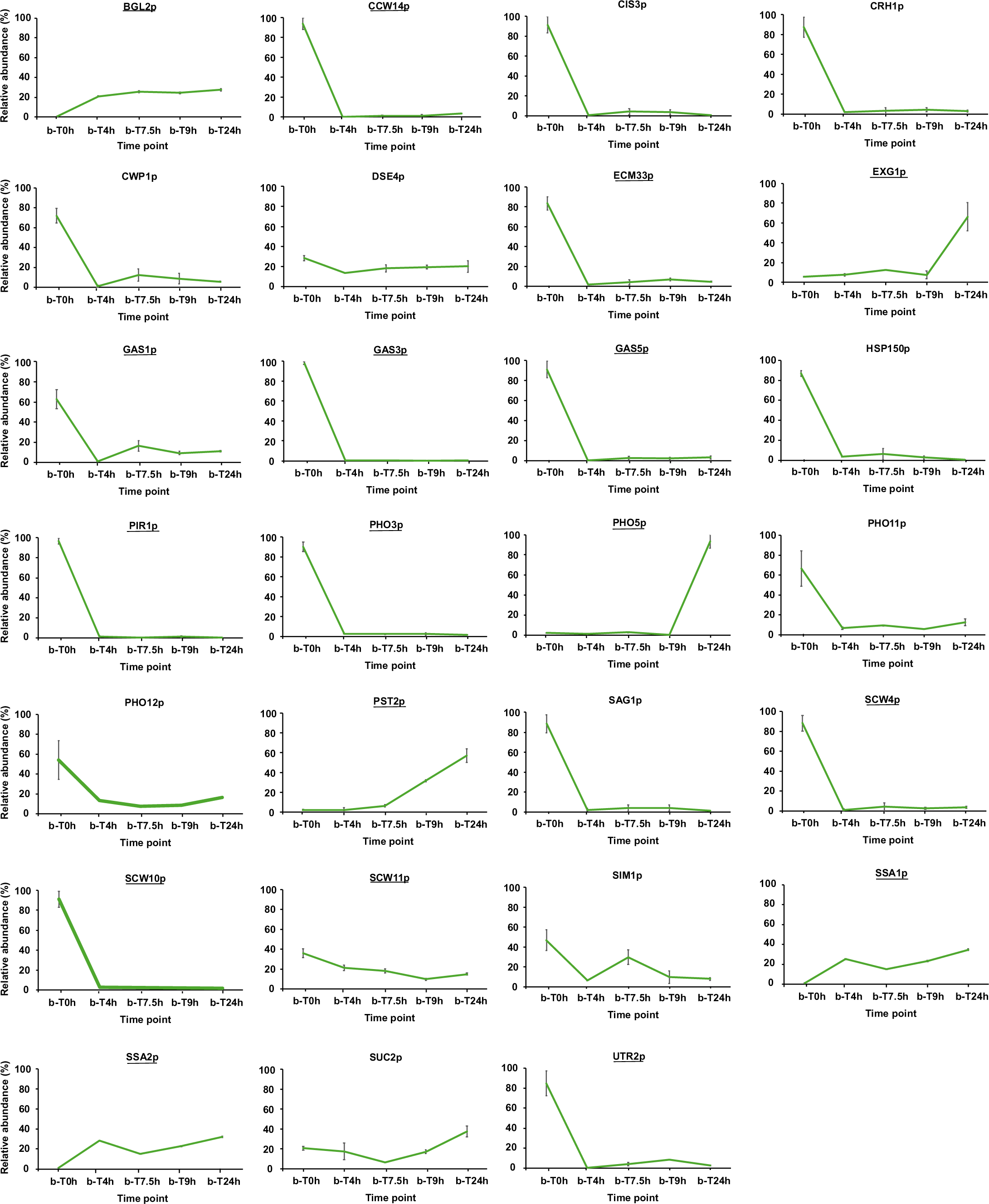

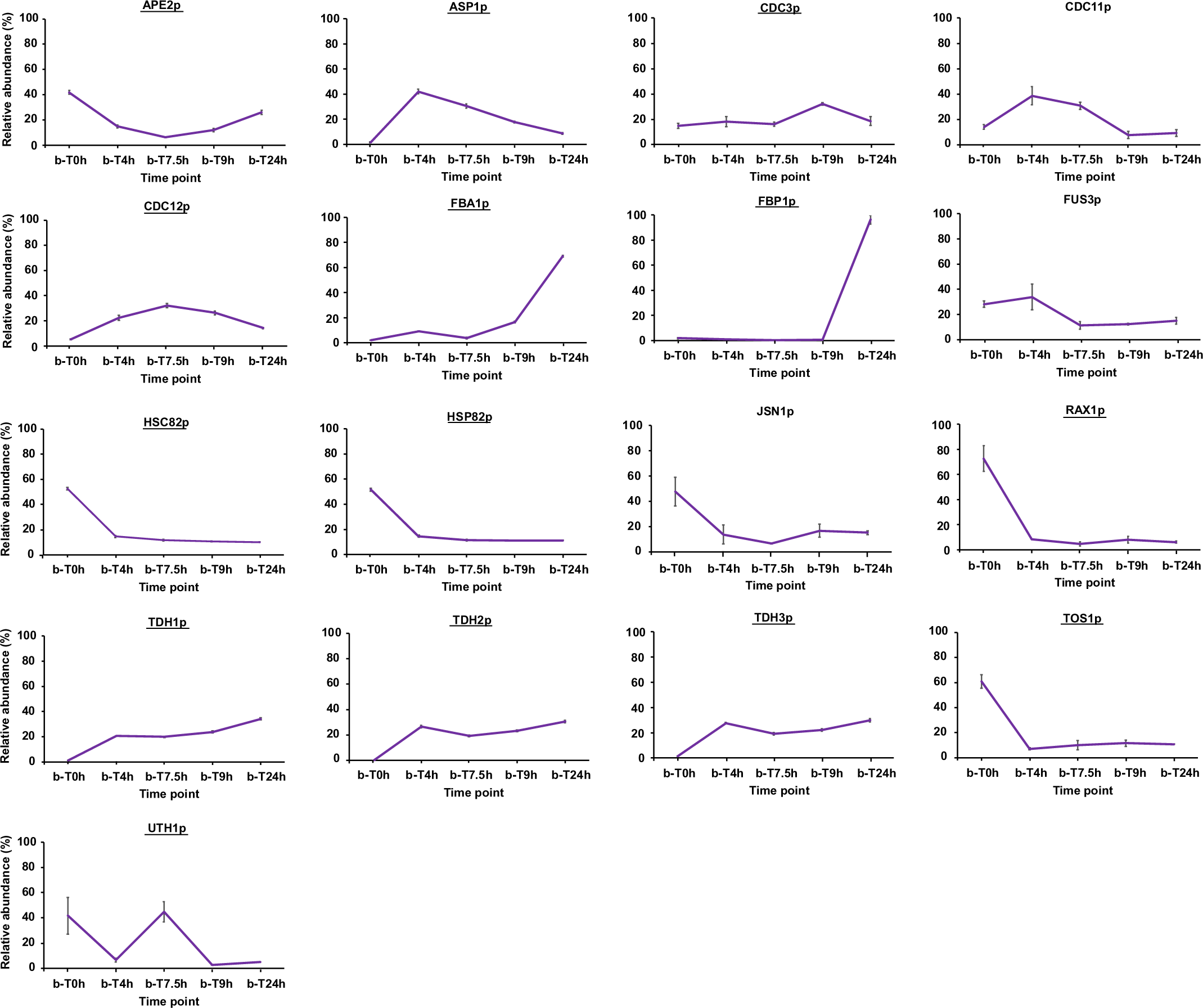

As shown in Fig. 4, the kinetic expression profiles of several CWPs change over time during batch cultivation. Most CWPs display their highest levels at b-T0h and reach their lowest expression at b-T4h. This aligns with the fact that these proteins are needed for transition stress at b-T0h and their expression is downregulated when cell wall is flexible during early exponential phase. Among these proteins, some that are linked to stress adaptation as well as cell wall remodeling and reinforcement show a slight increase in their expression over time during the diauxic shift and then decrease at the end of fermentation (Cis3p, Crh1p, Cwp1p, Ecm33p, Gas1p, Hsp150p, Sag1p, Scw4p, Utr2p), while others display a slight increase at the end of fermentation as Pho3p, Pho11p and Pho12p, consistent with their involvement in nutrient signaling during depletion at the end of the culture. The remaining proteins show a relatively flat kinetic profile throughout the culture (Gas3p, Gas5p, Pho3p, Pir1p, Scw10p), consistent with their role as YCW maintenance proteins that do not depend strongly on nutrient or stress pathways. Scw11p displayed a progressive decrease in abundance throughout fermentation, reaching its lowest level at the late diauxic shift. This decline corresponds to the reduced need for cell wall remodeling as cell division slows and cells transition toward a quiescent, nutrient-limited state. Toward the end of fermentation, Scw11p levels showed a modest increase, suggesting limited resumption of cell wall turnover associated with stationary-phase maintenance and long-term stress adaptation. Pho12p exhibited a trend comparable to Scw11p, displaying an expression pattern that was inversely related to Pho5p. Sim1p expression shows a complex, multiphasic expression pattern, peaking when yeast cells undergo major physiological transitions, as initiation and the early diauxic shift. This suggests a role in cell wall remodeling and adaptation to metabolic stress. Its decline during exponential growth and toward the end of fermentation reflects reduced need for such remodeling when cells are either rapidly proliferating or entering quiescence.

Bgl2p, Ssa1p, and Ssa2p follow similar trends: their expression is minimal at b-T0h, increases during the batch process, and then slightly vary for the rest of the culture. This expression pattern might be explained by the low activity of quiescent cells at b-T0h, where cells are not experiencing a strong growth-related cell wall expansion. Upregulation during the batch culture aligns with the need for increased glucan remodeling and protein folding demand, as cells are actively dividing, and they stabilize once a stable physiological state is reached.

In contrast, Exg1p, Pst2p, Suc2p and Pho5p exhibit fluctuations over time, reaching their maximum expression at the end of fermentation (b-T24h). The late-culture upregulation suggests that these proteins play a protective and adaptive role during nutrient depletion, helping cells to remodel and maintain the YCW under stress (Exg1p, Pst2p) and optimize phosphate acquisition (Pho5p) and carbon metabolism (Suc2p) when environmental nutrients are low, to survive prolonged culture conditions. This reflects the transition from growth-focused metabolism to survival and stress-adaptation mode.

Dse4p maintained low, relatively constant abundance throughout the batch culture, reflecting its role as a basal cell wall protein with a specialized, non-dominant function in these conditions.

CWPs that did not show significant variation across the culture according to the ANOVA test displayed relatively higher variability, as observed for Cis3p, Hsp150p, Cwp1p, Crh1p, Dse4p, Pho11p, Pho12p, Sag1p, Sim1p, and Suc2p. These proteins represent either structural CWPs heavily *O-*glycosylated that are integrated into the cell wall matrix or CWPs present at low abundance. The same analysis was carried out for the CWPs that can be found in other organelles (Fig. 5). Apart from the slight variation between replicates for each of these proteins (except for Uth1p, Jsn1p, Rax1p), we can clearly see that their expression is time-dependent during the batch culture. The expression of Tdh1p, Tdh2p and Tdh3p show the same trend as Ssa1p and Ssap2, with a minimal value at b-T0h where the cells are quiescent and is increased during the batch culture while remaining relatively constant all along, showing their implication in actively dividing cells. The expression of Hsp82p, Hsc82p, Rax1p and Tos1p showed the opposite trends, where their expression is maximal at b-T0h during the quiescence followed by a decrease and then stable levels during the rest of the batch culture. This pattern is consistent with early-phase stress adaptation, cell cycle/morphogenesis transitions, and the shift from quiescence to exponential growth.

Asp1p, Cdc11p, and Fus3p show a peak at the exponential phase (b-T4h), followed by a later decrease during the culture. This dynamic expression profile might refer to their role in supporting active division and growth-related signaling. As conditions worsen and growth slows, their expression naturally decreases as part of the transition toward stationary-phase physiology. They are also of low abundance compared to the other CWPs that can be found in other organelles. The other septins Cdc12p and Cdc3p expression profile show a peak at the early diauxic phase (b-T7.5h) and the late diauxic phase (b-T9h) respectively. Indeed, the diauxic shift requires reorganization and reinforcement of septin structures, with Cdc12p upregulated during early cytoskeletal remodeling and Cdc3p rising later to support long-term septin stability as cells approach stationary phase. Nevertheless, Uth1p, member of SUN family, showed a highly dynamic expression during the batch culture, peaking at the initiation (b-T0h) and the early diauxic phase (b-T7.5h), likely reflecting its regulation by multiple stress and metabolic pathways. Its roles in mitochondrial function, oxidative stress response, and cell wall remodeling make its abundance particularly sensitive to the changing physiological conditions encountered throughout fermentation.

Jsn1p and Apep displayed a similar expression profile across the culture with a maximum at b-T0h and a slight decrease during the early diauxic shift. This mild reduction likely reflects the transient global downregulation of translation and metabolic reorganization that occurs when cells transition from fermentation to respiration. Because both proteins function in general growth and metabolic maintenance rather than carbon source–regulated pathways, their abundance remains largely constant throughout the batch culture.

Fba1p and Fbp1p showed slightly variable expression in the early stages of batch culture, reflecting the metabolic transition from stationary-phase carryover to active fermentation. Their abundances increased markedly toward the end of the culture, reaching a maximum at b-T24h, consistent with glucose depletion and the induction of gluconeogenesis and respiratory metabolism. The strong late-stage increase in Fbp1p corresponds to its role as a glucose-repressed, starvation-induced enzyme, while Fba1p remains required for both glycolytic and gluconeogenic fluxes under nutrient-limited conditions.

### Comparison with Published Proteomic Analyses of Dynamics in Batch Culture

To assess the robustness and biological coherence of the temporal patterns observed in our dataset, we compared our results with two published proteomic studies that examined protein dynamics in *S. cerevisiae* cultured in batch mode.(den Ridder et al., 2023; Murphy et al., 2015) Both studies used TMT-based quantitative proteomics and sampled multiple growth phases, providing an appropriate benchmark for evaluating CWP expression trends.

Den Ridder *et al*. profiled two strains, CEN.PK113-7D (wild type) and an engineered minimal glycolysis mutant (MG), grown under aerobic and anaerobic conditions in controlled bioreactors.(den Ridder et al., 2023) Using a TMT 10-plex workflow, they quantified 36 CWPs, fewer than those detected in our study (supplemental Table S4). Their dataset revealed three major classes of CWP behavior (supplemental Fig. S6): (i) proteins whose abundance was primarily condition-dependent depending on oxygen availability (e.g., Scw4p, Gas3p, Ssa1p, Bgl2p, Pst2p, Suc2p and Exg1p); (ii) proteins with strain-specific expression (e.g., Hsp82p, Tdh3p); and (iii) proteins influenced by both strain and growth condition, including several strictly cell wall–localized proteins such as Gas1p, Ssa2p, Cis3p, Utr2p, Ygp1p, Pst1p, Gas5p, Ecm33p. A subset of CWPs was detected exclusively in particular strain - condition combinations : for example, Yps1p, Sim1p, Pho3p, Pho5p, Pho11p, and Pho12p appeared only in the MG strain under anaerobic conditions. Many CWPs showed little temporal variation across growth phases, while correlation analyses identified predominantly positive associations among strictly cell wall - localized proteins (supplemental Fig. S7). Overall, the study demonstrated condition- and strain-dependent regulation of CWP abundance, consistent with the dynamic shifts observed in our dataset.

Murphy *<u>et al.</u>* investigated proteomic remodeling during the diauxic shift in two wild-type strains (BY4742 and DBY7286) and a Hap2p-deficient mutant cultured aerobically in YPD in flasks.(Murphy et al., 2015) Their TMT 10-plex coupled to 2D-LC MS³ workflow quantified 56 CWPs in BY4742 and 51 CWPs in DBY7286, including several proteins not detected in our dataset (supplemental Table S5). In BY4742, CWPs segregated into Hap2-dependent and Hap2-independent groups (supplemental Fig. S8. A and B). Hap2-dependent proteins exhibited either continuous accumulation throughout culture (Ccw14p, Pst2p, Ssa1p, and Ygp1p) or a mid-culture peak followed by a decline (Cis3p, Cts1p, Cwp1p, Dse4p, Ecm33p, Exg1p, Pho5p, Pho11p, Pry3p, Pst1p, and Scw11p), while certain proteins (e.g., Pho3p) remained stable. Hap2p deletion strongly upregulated proteins such as Plb1p and Tdh1p (supplemental Fig. S9). In the DBY7286 strain, strictly cell wall - localized CWPs displayed four major temporal profiles, including mid-culture peaks (as for: Aga2p, Bar1p, Cis3p, Ecm33p, Exg2p, Gas1p, Gas3p, Scw4p, Scw10p, Scw11p, Sim1p, Pho3p, Pho11p), gradual increases (as for: Ath1p, Gas5p, Pst1p, Pst2p, Ssa1p, Ygp1p), or progressive decreases (as for: Bgl2p, Cts2p, YJL171Cp, Ssa2p, Suc2p) or even variable complex trends (as for: Cwp1p, Exg1p and Ccw14p) (supplemental Fig. S10). CWPs with multi-organelle localization exhibited six additional temporal patterns, reflecting the metabolic shift associated with glucose depletion and the onset of respiration (supplemental Fig. S11).

Comparison of CWP sequence coverage across the studies showed that our dataset generally yielded higher values, particularly for CWPs strictly localized to the YCW and for TDH proteins (supplemental Fig. S12 and supplemental Table S6). This likely reflects the sample preparation strategy used in our study, which specifically enriched the YCW fraction. In contrast, CWPs that may localize to other organelles exhibited higher sequence coverage in the study by Murphy *et al*., likely due to the extensive fractionation and separation methods applied in their workflow (supplemental Fig. S12 and supplemental Table S6).

Across both external datasets, CWPs consistently showed dynamic phase-dependent expression, paralleling and complementing the patterns observed in our study. Moreover, a genotype-specific expression was also shown in their studies. While differences in strain backgrounds, culture conditions (bioreactor vs. flask), and analytical workflows likely contributed to variation in detected proteins and temporal profiles, the overarching agreement across studies supports the biological robustness of our findings. Together, these comparisons reinforce the conclusion that CWP remodeling is a conserved and reproducible feature of *S. cerevisiae* growth in batch culture.

## Discussion

This study aimed to investigate the dynamic change of YCW proteins expression during a batch culture. To this end, we used firmly regulated bioreactors and carried out the experiments in biological triplicates with well-established sample preparation methods, yielding a high reproducibility and precision. Although several previous studies have examined the temporal dynamics of the *Saccharomyces cerevisiae* proteome under various culture conditions (Costenoble et al., 2011; de Groot et al., 2007; den Ridder et al., 2023; Helbig et al., 2009; Murphy et al., 2015), none has specifically focused on the proteome dynamics at the level of the cell wall. Moreover, in the early 2000s, Klis and colleagues conducted proteomic analyses to identify and quantify CWPs in *Saccharomyces cerevisiae* strains grown in flask-based batch cultures, employing both label-free and label-based approaches (Yin et al., 2005). In their work, Klis and co-workers identified and quantified 19 CWPs that were consistently detected across all growth phases following proteolytic digestion directly or subsequently to chemical or enzymatic fractionation of YCW isolated by mechanical disruption, except for Pir3p, which was uniquely observed during the stationary phase of wild type and mutant strains of FY833 cultivated in batch mode in YPD medium within flasks (Yin et al., 2005). However, mass spectrometry technologies have since advanced substantially, allowing for far more comprehensive and sensitive proteome mapping. These developments underscore the importance of revisiting such analyses using modern instrumentation and strictly controlled conditions, as enabled by bioreactor systems.

The quantification of approximately 1,500 proteins in our study highlights the depth of our proteomic analysis while suggesting that additional optimization of YCW preparation could enhance specificity (Yammine et al., 2022). This number is comparable to that reported by den Ridder *et al*. (den Ridder et al., 2023), who analyzed total cell lysates without prior fractionation or enrichment. Nevertheless, the relative abundance of CWPs was markedly higher in our dataset as well as the sequence coverage of CWPs, demonstrating that our protocol effectively enriched the yeast cell wall fraction. In Murphy and collaborators’ work, a significantly higher number of proteins exceeding 3,500 was obtained, due to extensive fractionation using high pH reverse phase and bidimensional LC separation of peptides following digestion of the total cell lysates of yeast cells (Murphy et al., 2015).

Differential proteomics using univariate and multivariate statistical analyses coupled to functional enrichment analyses revealed how carbon source limitation affects the metabolism in general and the CWPs expression in particular. In the exponential phase, the proteomes are enriched with ribosomal proteins and proteins involved in cellular growth and protein synthesis, and less enriched in CWPs. During the exponential phase, cells prioritize growth and division, resulting in a relatively flexible cell wall optimized for rapid expansion. CWPs expressed at this stage included enzymes responsible for glucan assembly and restructuring involved in cell wall biosynthesis and remodeling (i.e. Bgl2p and Scw11p, as well as Asp1p, Cdc11p, Cdc12p and Fus3p in our dataset). Proteomic comparison across growth phases revealed that the diauxic shift induces the most pronounced proteomic rearrangement in *S. cerevisiae*. This transition, triggered by glucose depletion and the linear increase in ethanol concentration, shifts metabolism from respiro-fermentative to fully respiratory, increasing proteins related to respiration, fatty acid metabolism, and energy production while decreasing those involved in protein synthesis, due to major transcriptional reprogramming. These changes reflect very limited growth on ethanol and a reorganization toward survival and stress resistance, affecting the cell wall. Mitochondrial proteins increased as glycolytic enzymes declined, consistent with enhanced respiratory activity. The metabolic states of yeast cells grown in bioreactor differ significantly between the pre- and post-diauxic phases. The upregulation of stress-response proteins, such as heat shock proteins, further supports improved robustness under nutrient limitation, and start appearing in the wall (i.e. Hsp150p, Cis3p and Cwp1p in our dataset), along with enzymes that help their incorporation and cell wall reorganization (i.e. Cdc3p, Gas1p, Sim1p, Uth1p and Utr2p). Therefore, cell wall begins to strengthen in preparation for harsher conditions.

Notably, the reduction in glycolytic enzymes and ribosomal proteins during the stationary phase may delay rapid regrowth upon re-exposure to glucose, relevant for optimizing industrial fermentation processes. These outcomes support and extend findings from previous investigations (den Ridder et al., 2023; Fuge et al., 1994; Murphy et al., 2015; Valcourt et al., 2012; Werner-Washburne et al., 1996).

Quiescence is a reversible, non-dividing state that yeast cells adopt under nutrient limitation, particularly carbon or nitrogen depletion. In this state, quiescent cells remain metabolically active but show minimal proliferation, enhanced stress resistance, and increased longevity. In our study, proteins associated with the “cell periphery” and “plasma membrane” were more abundant, reflecting cell wall and membrane remodeling that strengthens structural integrity and stress tolerance, hallmarks of quiescent physiology. In addition to the incorporation of stress resistance proteins (i.e. Pir proteins), cross-linking of glucans and chitin is increased for structural rigidity. The induction of these proteins seems to be associated with glucose starvation and oxidative stress (Ribeiro et al., 2022).

Both b-T0h and b-T24h correspond to stationary phases marked by growth arrest and entry into quiescence. In the seed culture, the medium is not aerated despite agitation, resulting in very limited oxygen transfer and near-anaerobic growth. Metabolism is therefore predominantly fermentative, and pH is not controlled. Consequently, growth conditions differ markedly from those of batch cultures in a bioreactor, irrespective of the sampling time, which likely explains the atypical proteomic profiles observed. However, the proteomic profile at b-T0h likely reflects additional stress responses due to uncontrolled pH and oxygen conditions in flask-based batch culture, with increased expression of enzymes as Ape2p, Crh1p, Gas3p, Gas5p, Scw4p, Pho3p, Pho12p and Dse4p, structural proteins as Ccw14p, Ecm33p, Pir1p, Sag1p, heat shock proteins as Hsc82p and Hsp82p as well as other proteins with unknow functions like Jsn1p, Rax1p and Tos1p. On the other hand, b-T24h represents a more stable quiescent state under bioreactor regulation, with upregulation of enzymes as Exg1p, Pho5p, Pst2p, Suc2p along with Fba1p and Fbp1p for gluconeogenesis. These findings emphasize cell wall remodeling as a key adaptive mechanism in the establishment and maintenance of quiescence in *S. cerevisiae*. Functionally, YCW remodeling enhances resistance to various stresses, including oxidative damage and osmotic pressure. It also enables yeast to enter and sustain dormancy, survive desiccation, and withstand antimicrobial agents (Jin et al., 2022; Sanz et al., 2017). Throughout the bioreactor culture, Tdh1p, Tdh2p, and Tdh3p, along with the molecular chaperones Ssa1p and Ssa2p, exhibited consistently elevated expression compared with the initial time point (b-T0h). The sustained upregulation of the TDHp isoforms, key glycolytic enzymes, suggests ongoing energy metabolism to meet cellular demands during different growth phases, while persistent Ssa1p and Ssa2p expression indicates continuous engagement of protein folding and quality control mechanisms. Together, these patterns reflect a coordinated strategy whereby yeast maintains metabolic activity and stress-resilience machinery to support adaptation to dynamic environmental and nutritional conditions in batch culture. Such sustained expression may also underpin the capacity of cells to remodel the cell wall, manage oxidative or heat stress, and prepare for stationary-phase survival.

The dynamic distribution of transcription factors across growth phases underscores the complexity of the regulatory networks driving yeast adaptation during batch culture. The predominance of shared transcription factors suggests a core regulatory program balancing protein synthesis and stress adaptation, while the small subset of phase-specific regulators likely fine-tunes responses to environmental and metabolic shifts. The enrichment of upregulated proteins at the cell periphery and mitochondria points to enhanced cell wall remodeling and energy metabolism during adaptation, whereas the repression of nucleosomal proteins may reflect a transition toward maintenance and stress resistance rather than transcription and proliferation. Notably, the uncharacterized transcription factors YKL222Cp and YPR196Wp appear to induce specific pathways that need to be further investigated.

By benchmarking our results to previous studies, we realized that the differences observed compared with the studies by Murphy (Murphy et al., 2015) and den Ridder (den Ridder et al., 2023) likely reflect variations in experimental design, including sample preparation (total protein lysates versus cell wall-enriched fractions), strain background, culture conditions, and proteomic analysis strategies. Such methodological distinctions can markedly influence protein extraction efficiency, detection sensitivity, and quantification, ultimately shaping the composition and interpretation of the reported cell wall proteomes. Our dataset revealed several CWPs not reported in earlier studies, demonstrating the strong detection capacity of our workflow. The fact that each prior study also identified unique proteins highlights the added value of integrating multiple datasets to achieve a more comprehensive view of the cell wall proteome.

Comparison of exclusively localized CWPs across studies revealed that each dataset captured distinct components of the yeast cell wall proteome, underscoring the complementary nature of these analyses. Notably, our dataset uniquely identified Sag1p and Pir1p, that are two proteins central to adhesion, structural reinforcement, and stress resilience. This demonstrates the strong detection depth and biological relevance of our workflow. Other studies similarly reported proteins absent from our analysis, reflecting condition- and strain-specific expression; for example, Murphy *et al*. identified CWPs such as Kre9p, Dse4p, and Pry3p in S288c-derived strains, as well as Aga2p, Bar1p, and Ath1p in DBY7286, highlighting pathways linked to cytokinesis, mating, and metabolic adaptation (Murphy et al., 2015). A few CWPs not detected in our dataset (Ygp1p, Plb2p, Pst1p) likely represent proteins expressed only under specific starvation or stress conditions, reinforcing the context-dependence of CWP composition. Similarly, proteins detected in other studies - such as Rax2p, Bud4p, Kss1p, and GPI-anchored proteases Yps1p and Mkc7p - reflect dynamic remodeling processes tied to morphogenesis, polarity, and signaling. Importantly, Murphy *et al*. further distinguished Hap2p-dependent CWPs, showing that regulatory networks selectively modulate subsets of structural and stress-responsive proteins, while the study by den Ridder *et al*. revealed oxygen-dependent CWP remodeling not captured under our culture conditions (den Ridder et al., 2023). Overall, the benchmarking analysis highlights the robustness of our dataset, which captured key structural and adaptive CWPs, including unique elements not previously reported, while also illustrating how differences in strain background, growth environment, and proteomic workflows shape the detectable cell wall proteome. Future work using refined YCW purification will improve the detection of low-abundance CWPs (Yammine et al., 2022). This approach will be implemented in forthcoming experiments. Moreover, we will use this strategy to explore how factors such as temperature and pH further influence cell wall composition and adaptation. Time-course proteomics of *S. cerevisiae* in batch culture reveals dynamic cell wall remodeling as a tightly coordinated adaptive response. These changes align with metabolic shifts and environmental challenges, enabling the cell wall to actively support yeast survival across growth phases, with implications for both fundamental biology and industrial applications.

## Abbreviations

ANOVA: Analysis of Variance
CWP: Cell Wall Protein
EDTA: 2,2’,2’’,2’’’-(Ethane-1,2-diyldinitrilo)tetraacetic acid
eFASP: enhanced filter-assisted sample preparation
FDR: false Discovery Rate
GO: Gene Ontology
GPI: Glycosylphosphatidylinositol
H_2_SO_4_: Sulfuric acid
HPLC: High Performance Liquid Chromatography
KEGG: Kyoto Encyclopedia of Genes and Genomes
(2D-)LC: (bidimensional) Liquid Chromatography
MG: Minimal Glycolysis
NaCl: Sodium chloride
NaOH: Sodium hydroxide
OD: Optical density
PCA: Principal Component Analysis
SDS: Sodium dodecyl sulfate
SGD: Saccharomyces Genome Database
Tris-HCl: 2-Amino-2-(hydroxymethyl)propane-1,3-diol, chlorhydrate
YCW: Yeast Cell Wall
YPD: Yeast Extract – Peptone – Dextrose

## Data availability

All raw and processed proteomics data supporting the findings of this study have been deposited in the PRIDE repository under the dataset identifier PXD071725 and are publicly accessible.

## Supplemental data

Supplemental materials for this study are available online and include nine figures (in PDF file) and five tables (Excel format). The supplemental figures (Fig. S1-S12) provide additional information about the growth in batch culture mode in bioreactors, as well as visualization of CWP expression patterns, temporal dynamics, and comparative analyses of CWP expression and sequence coverage across strains and conditions in other studies, supporting the interpretations in Figures 3–5 of the main text. The supplemental tables (S1–S6) contain detailed quantitative proteomics data, lists of differentially expressed proteins, functional annotations and sequence coverage information that underpin the analyses discussed throughout the manuscript.

## Acknowledgments

The authors gratefully acknowledge Laura Mena, Pauline Bellanger, Alexis Lavogiez, and Nicolas Klemesiak for their valuable technical assistance in handling and monitoring yeast fermentations in bioreactors.

## Funding and Additional Information

The Ph.D. funding of M.Y. was received from a joint grant awarded by I-Site Université Lille Nord-Europe, Région Hauts-de-France, and Lesaffre International. The mass spectrometry analyses were carried out on the Top_Omics platform, supported by funding from the EU European Regional Development Fund (ERDF), the Région Hauts-de-France, the CNRS, and the University of Lille. The platform is recognized as an IBISA (Infrastructures en Biologie, Santé et Agronomie) facility and is a member of two national infrastructures ProFI (FR2048 CNRS ) and Infranalytics (FR2048 CNRS). Yeast fermentations in batch mode were performed at the Lesaffre International Research and Development facilities. The graphical abstract was created with BioRender.com.

## Author contributions

M. Y. , C. R. , E. P. and I. M. conceptualization; A. P. , M. Y. , C. R. , E. P. , F. B. , I. M. and S. F. methodology; C. R. , E. P. and I. M. funding acquisition; M. Y. , C. R., F. B. and I. M. investigation; A. P. , M. Y. , C. R. , E. P. visualization; A. P. , M. Y. , C. R. , E. P. formal analysis; C. R. , I. M. supervision; M. Y. writing-original draft; A. P. , M. Y. , C. R. , E. P. , I. M. and writing review and editing.

## Conflict of interest

E.P. , A.P. , and I.M. are employed by Lesaffre International. The other co-authors declare no competing interests.

