## Supplemental information for "Time-Resolved Analysis of the Cell Wall Proteome in *Saccharomyces cerevisiae* S288c During Batch Fermentation"

**Table of contents**

| Content | Title | Page |
| --- | --- | --- |
| SI Figure 1. | ***S. cerevisiae* S288c growth in bioreactor batch culture.** | **S-3** |
| SI Table 1. | **Pearson correlation coefficients of proteomic profiles across biological replicates and growth phases during batch fermentations in bioreactor of S288c yeast.** | **S-4** |
| SI Figure 2. | **Functional enrichment analysis with KEGG pathways of differentially upregulated proteins in different time points during the batch fermentation in bioreactor of *S. cerevisiae* S288c yeast.** | **S-5** |
| SI Figure 3. | **Functional enrichment analysis with KEGG pathways of differentially downregulated proteins in different time points during the batch fermentation in bioreactor of *S. cerevisiae* S288c yeast.** | **S-6** |
| SI Figure 4. | **Functional enrichment analysis of the target genes regulated by transcription factors specifically associated with either upregulated or downregulated proteins using STRING-DB.** | **S-7** |
| SI Figure 5. | **Spearman rank correlation of CWPs expression during the kinetics of batch culture of S288c strain in bioreactor** **from our dataset, using Morpheus software.** | **S-8** |
| SI Figure 6. | **Hierarchical clustering heatmap by Euclidean distance for Z-score normalized protein abundances established with Morpheus software, showing time point-specific expression signatures of CWPs in S. cerevisiae grown in batch mode with a rich medium in bioreactor from den Ridder *et al* dataset.** | **S-9** |
| SI Figure 7. | **Spearman rank correlation of CWPs expression during the kinetics of batch culture of** **CEN.PK113-7D wild-type and mutant strains in bioreactor from den Ridder *et al* study, using Morpheus software.** | **S-10** |
| SI Figure 8. | **Relative abundance changes of strictly cell wall - localized CWPs during batch culture of the wild-type *S. cerevisiae* BY4742 strain derived from S288c strain and its Hap2p-deletion mutant in 1 L flasks from Murphy *et al*.** | **S-11**  **S-12** |

| Content | Title | Page |
| --- | --- | --- |
| SI Figure 9. | **Relative abundance changes of CWPs that are also localized in other organelles during batch culture of the wild-type *S. cerevisiae* BY4742 strain derived from S288c and its Hap2p-deletion mutant in 1 L flasks from Murphy *et al*.** | **S-13** |
| SI Figure 10. | **Relative abundance changes of strictly cell wall – localized CWPs during batch culture of the wild-type *S. cerevisiae* DBY7286 strain in 1 L flasks for 33 h from Murphy *et al.*** | **S-14** |
| SI Figure 11. | **Relative abundance changes of strictly cell wall – localized CWPs during batch culture of the wild-type *S. cerevisiae* DBY7286 strain in 1 L flasks for 33 h from Murphy *et al.*** | **S-15** |
| SI Figure 12. | **Comparison of CWP sequence coverage (%) between our dataset and those reported by Murphy *et al.* (21) and den Ridder *et al.* (22).** | **S-16** |

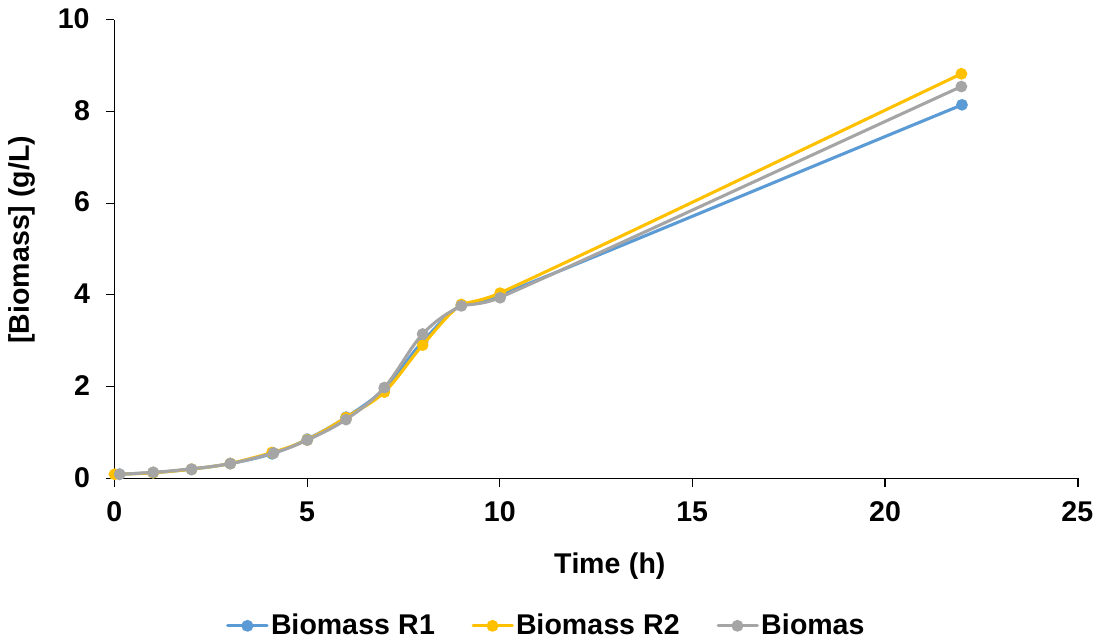
**A.**

**B.**

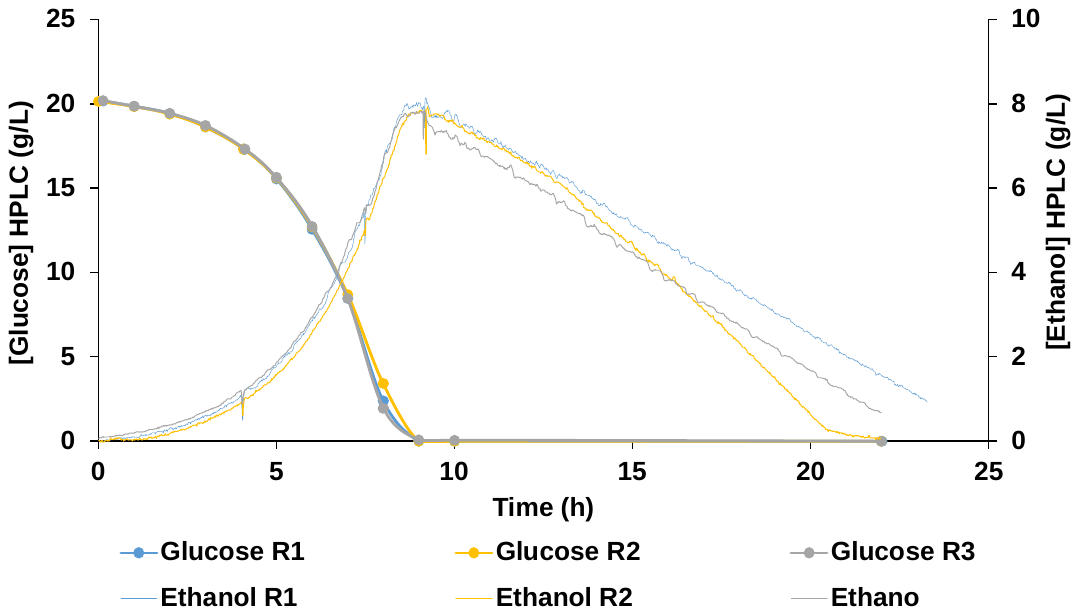

**SI Figure 1. S. cerevisiae S288c growth in bioreactor batch culture. A.** Growth curve of the three biological replicates representing the biomass produced during the cultures in g (Dry Weight)/L, calculated based on the optical density measurements of cells at 600 nm. **B.** Glucose consumption and Ethanol production for the three biological replicates determined by HPLC in the supernatant withdrawn during the ongoing of the cultures. R1, R2 and R3 refer to the number of the biological replicate.

**SI Table 1. Pearson correlation coefficients of proteomic profiles across biological replicates and growth phases during batch fermentations in bioreactor of S288c yeast.** To assess the data reproducibility and consistency between samples, column correlation function was applied using Perseus software v2.1.5.0. A color scale indicates the higher values (red) and the lower values (blue) of correlation coefficients.

| **b-T0h-R1** | **b-T0h-R2** | **b-T0h-R3** | **b-T4h-R1** | **b-T4h-R2** | **b-T4h-R3** | **b-T7.5h-R1** | **b-T7.5h-R2** | **b-T7.5h-R3** | **b-T9h-R1** | **b-T9h-R2** | **b-T9h-R3** | **b-T24h-R1** | **b-T24h-R2** | **b-T24h-R3** | **Name** |
| --- | --- | --- | --- | --- | --- | --- | --- | --- | --- | --- | --- | --- | --- | --- | --- |
| NaN | 0.86 | 0.89 | 0.51 | 0.54 | 0.55 | 0.52 | 0.51 | 0.57 | 0.43 | 0.41 | 0.50 | 0.27 | 0.29 | 0.27 | **b-T0h-R1** |
| 0.86 | NaN | 0.87 | 0.56 | 0.58 | 0.59 | 0.56 | 0.56 | 0.60 | 0.49 | 0.48 | 0.55 | 0.33 | 0.34 | 0.34 | **b-T0h-R2** |
| 0.89 | 0.87 | NaN | 0.53 | 0.55 | 0.55 | 0.53 | 0.53 | 0.58 | 0.46 | 0.44 | 0.52 | 0.30 | 0.32 | 0.30 | **b-T0h-R3** |
| 0.51 | 0.56 | 0.53 | NaN | 0.89 | 0.84 | 0.86 | 0.86 | 0.83 | 0.80 | 0.80 | 0.79 | 0.61 | 0.62 | 0.62 | **b-T4h-R1** |
| 0.54 | 0.58 | 0.55 | 0.89 | NaN | 0.88 | 0.86 | 0.86 | 0.85 | 0.81 | 0.82 | 0.82 | 0.62 | 0.63 | 0.63 | **b-T4h-R2** |
| 0.55 | 0.59 | 0.55 | 0.84 | 0.88 | NaN | 0.81 | 0.83 | 0.82 | 0.78 | 0.77 | 0.78 | 0.60 | 0.60 | 0.60 | **b-T4h-R3** |
| 0.52 | 0.56 | 0.53 | 0.86 | 0.86 | 0.81 | NaN | 0.88 | 0.85 | 0.84 | 0.84 | 0.83 | 0.64 | 0.66 | 0.65 | **b-T7.5h-R1** |
| 0.51 | 0.56 | 0.53 | 0.86 | 0.86 | 0.83 | 0.88 | NaN | 0.87 | 0.85 | 0.84 | 0.85 | 0.67 | 0.68 | 0.68 | **b-T7.5h-R2** |
| 0.57 | 0.60 | 0.58 | 0.83 | 0.85 | 0.82 | 0.85 | 0.87 | NaN | 0.80 | 0.79 | 0.84 | 0.62 | 0.62 | 0.61 | **b-T7.5h-R3** |
| 0.43 | 0.49 | 0.46 | 0.80 | 0.81 | 0.78 | 0.84 | 0.85 | 0.80 | NaN | 0.89 | 0.87 | 0.75 | 0.76 | 0.75 | **b-T9h-R1** |
| 0.41 | 0.48 | 0.44 | 0.80 | 0.82 | 0.77 | 0.84 | 0.84 | 0.79 | 0.89 | NaN | 0.86 | 0.76 | 0.75 | 0.76 | **b-T9h-R2** |
| 0.50 | 0.55 | 0.52 | 0.79 | 0.82 | 0.78 | 0.83 | 0.85 | 0.84 | 0.87 | 0.86 | NaN | 0.73 | 0.73 | 0.73 | **b-T9h-R3** |
| 0.27 | 0.33 | 0.30 | 0.61 | 0.62 | 0.60 | 0.64 | 0.67 | 0.62 | 0.75 | 0.76 | 0.73 | NaN | 0.90 | 0.91 | **b-T24h-R1** |
| 0.29 | 0.34 | 0.32 | 0.62 | 0.63 | 0.60 | 0.66 | 0.68 | 0.62 | 0.76 | 0.75 | 0.73 | 0.90 | NaN | 0.91 | **b-T24h-R2** |
| 0.27 | 0.34 | 0.30 | 0.62 | 0.63 | 0.60 | 0.65 | 0.68 | 0.61 | 0.75 | 0.76 | 0.73 | 0.91 | 0.91 | NaN | **b-T24h-R3** |

| 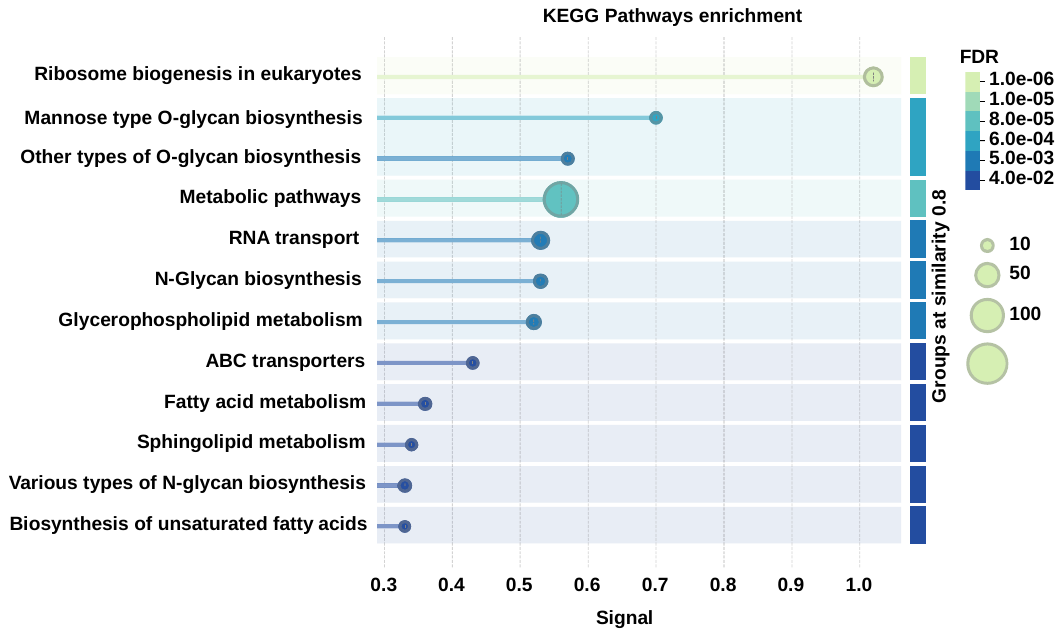**A.** | 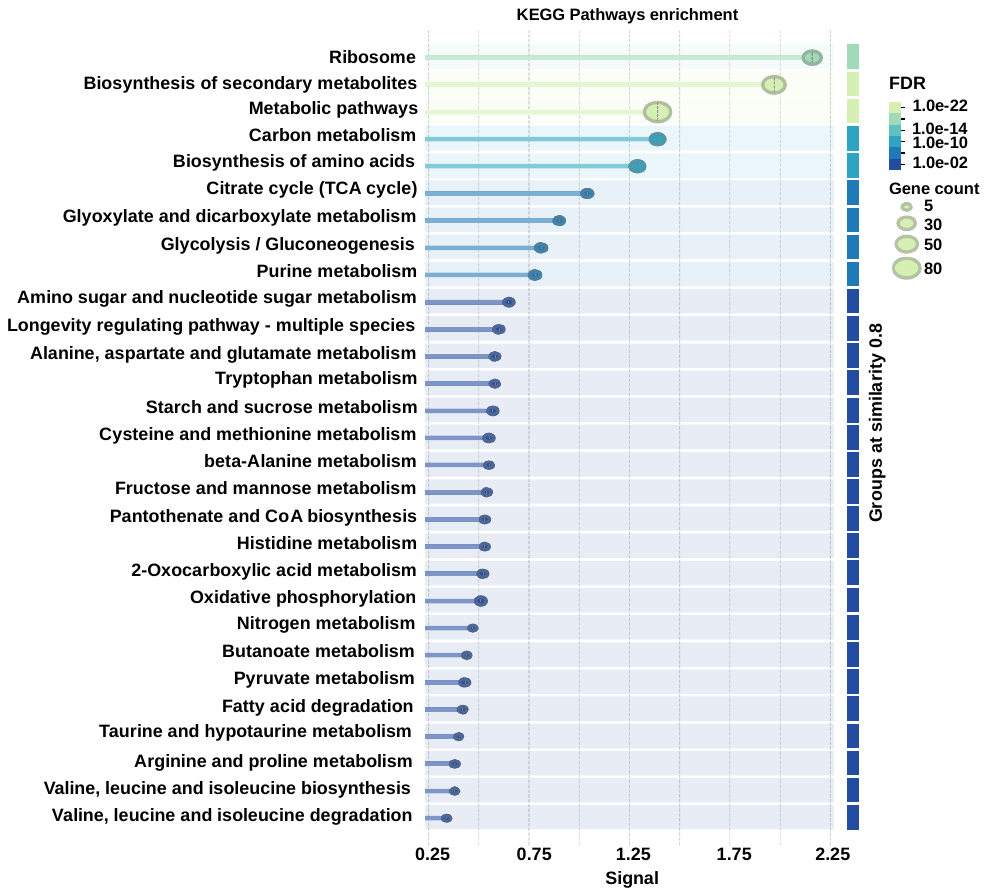**D.**  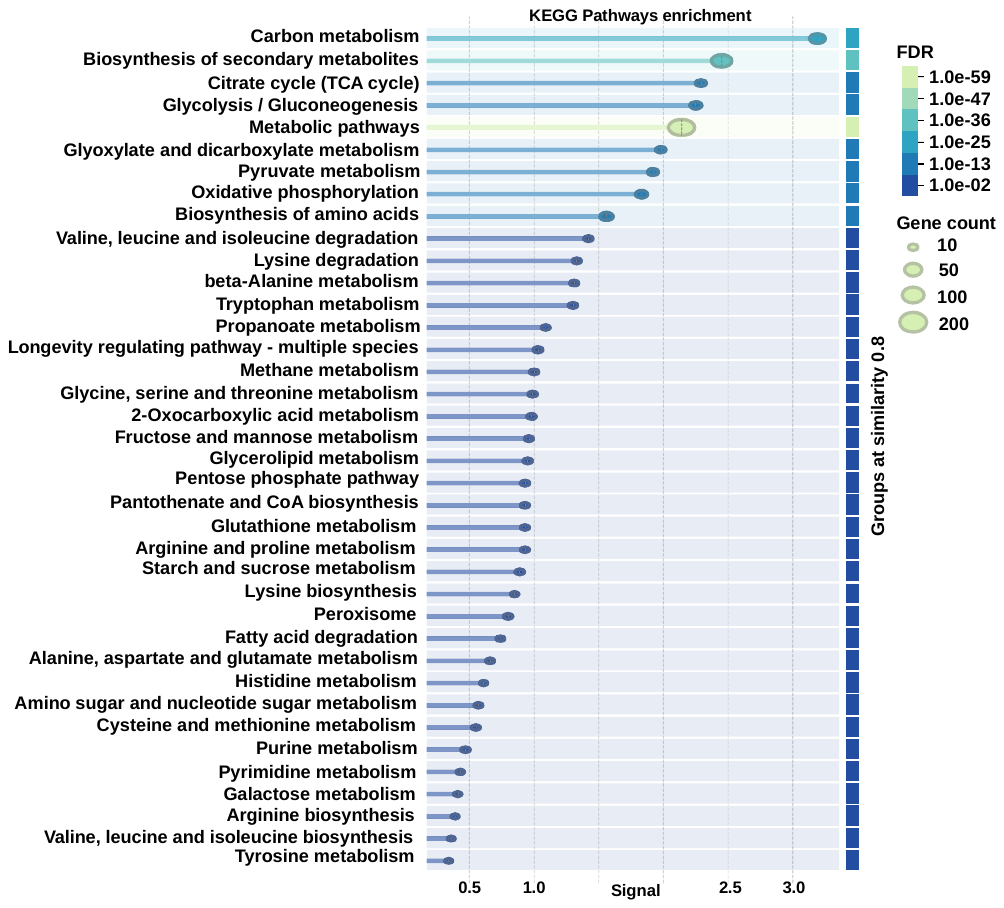**E.** |
| --- | --- |
| 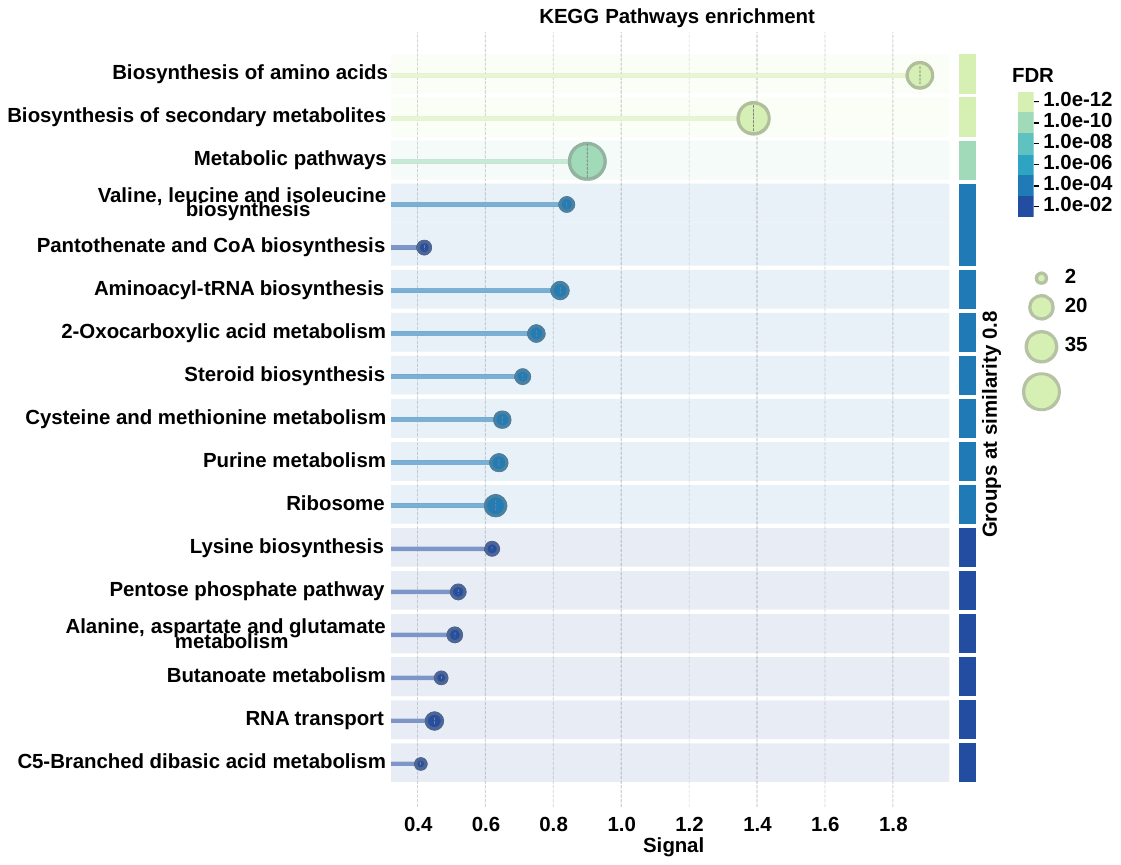**B.** |  |
| 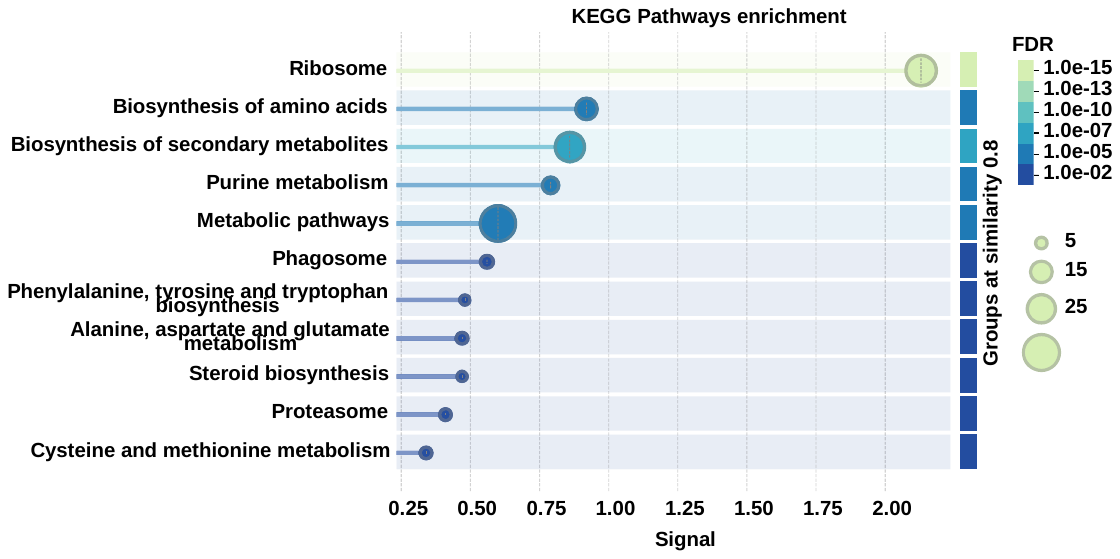**C.** |  |
| **SI Figure 2. Functional enrichment analysis with KEGG pathways of differentially upregulated proteins in different time points during the batch fermentation in bioreactor of *S. cerevisiae* S288c yeast.** The upregulated proteins that were commonly quantified in all comparisons between a specific time point (A. b-T0h; B. b-T4h, C. b-T7.5h, D. b-T9h, E. b-T24h) and the others were subjected to this analysis. | |

| **A.**  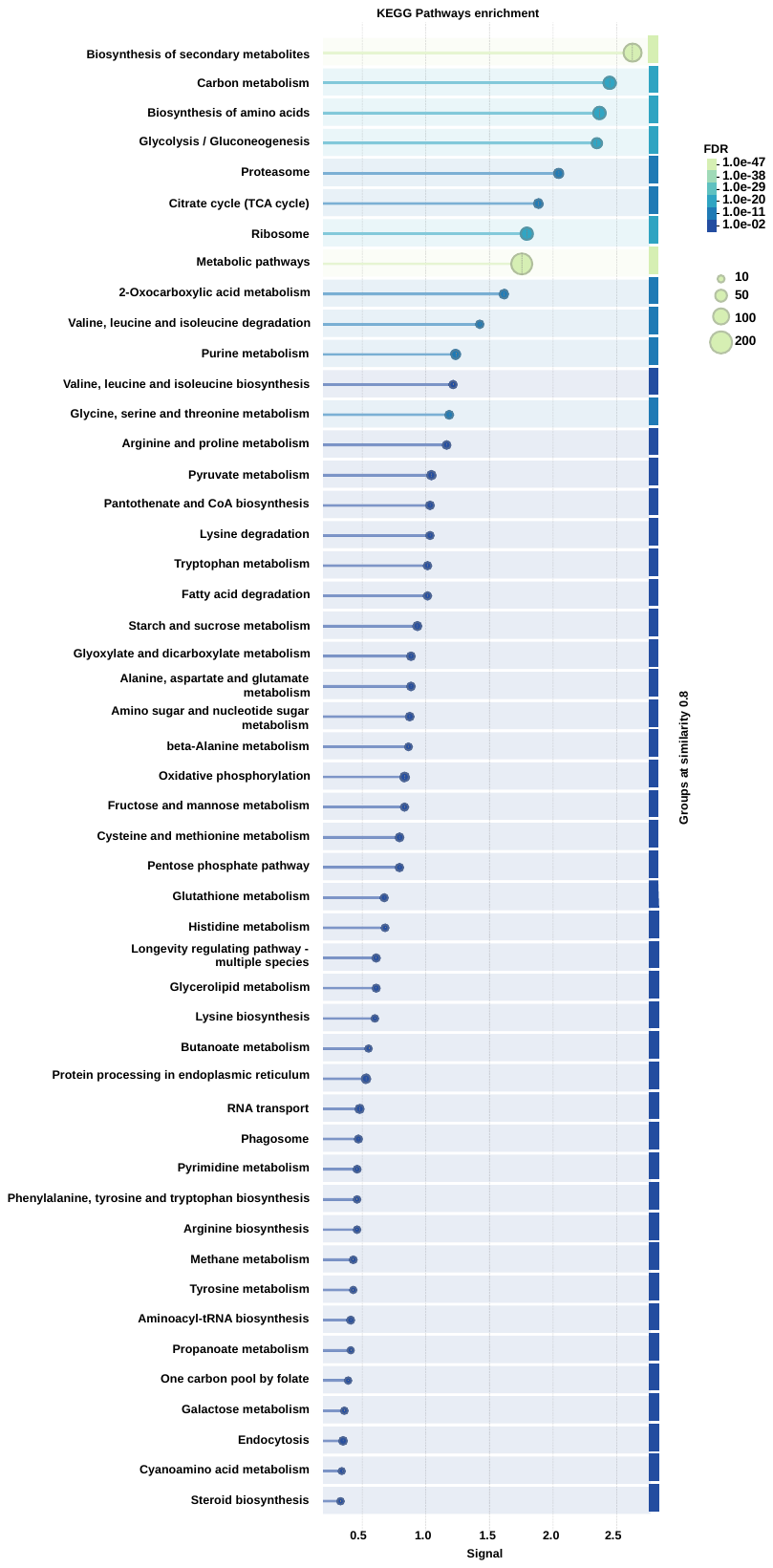 | 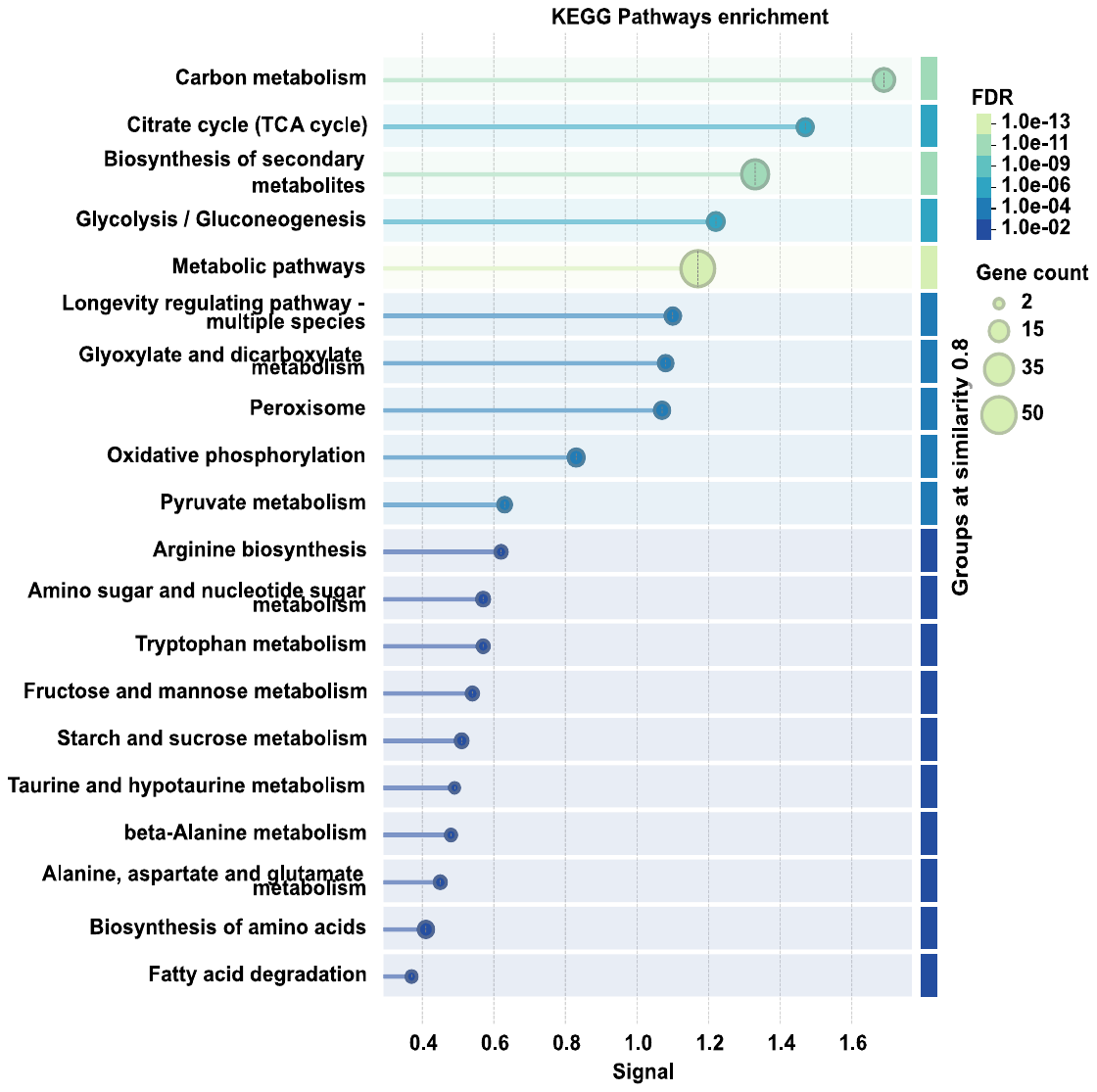**B.**  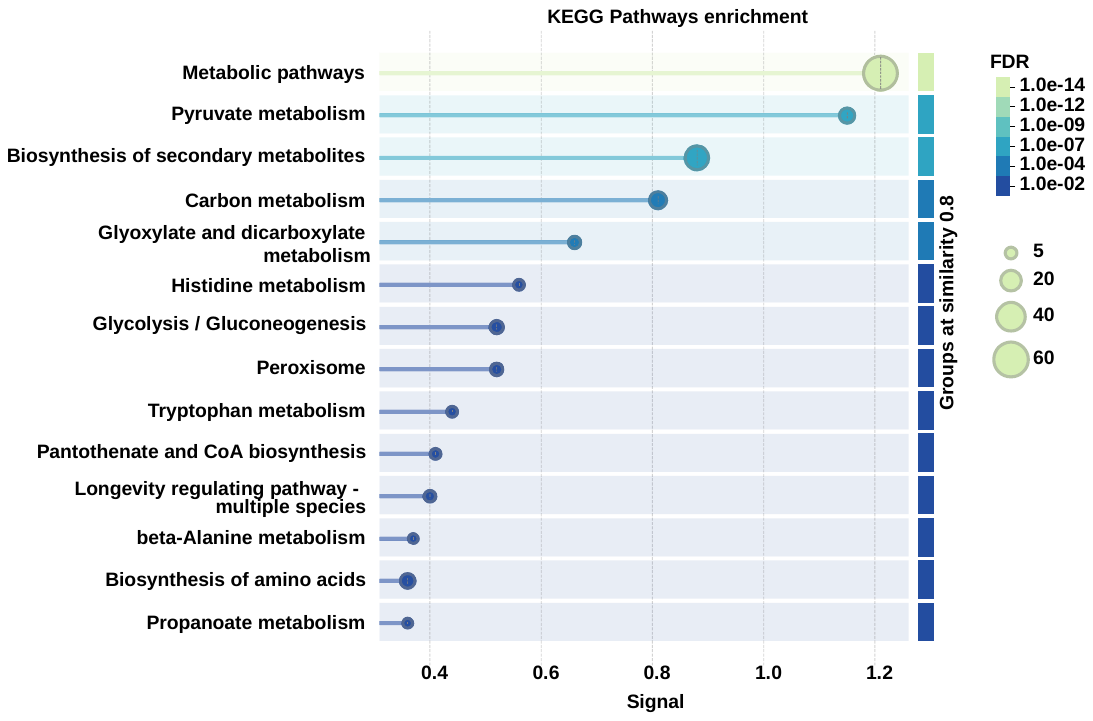**C.**  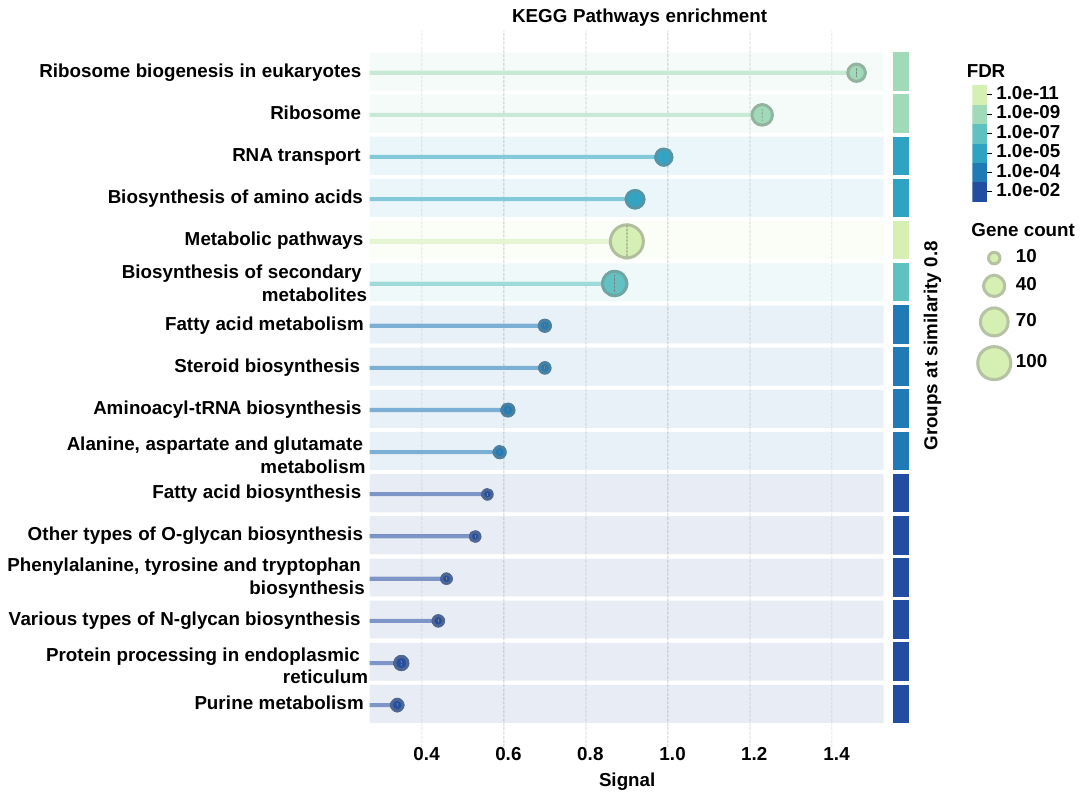**E.** |
| --- | --- |
| 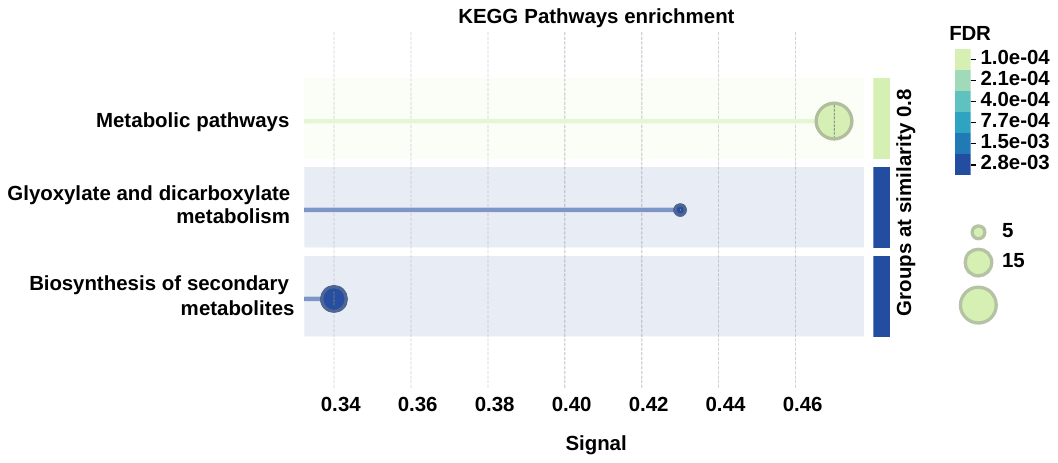**D.** |  |
| **SI Figure 3. Functional enrichment analysis with KEGG pathways of differentially downregulated proteins in different time points during the batch fermentation in bioreactor of *S. cerevisiae* S288c yeast.** The downregulated proteins that were commonly quantified in all comparisons between a specific time point (A. b-T0h; B. b-T4h, C. b-T7.5h, D. b-T9h, E. b-T24h) and the others were subjected to this analysis. | |

| **A.** | 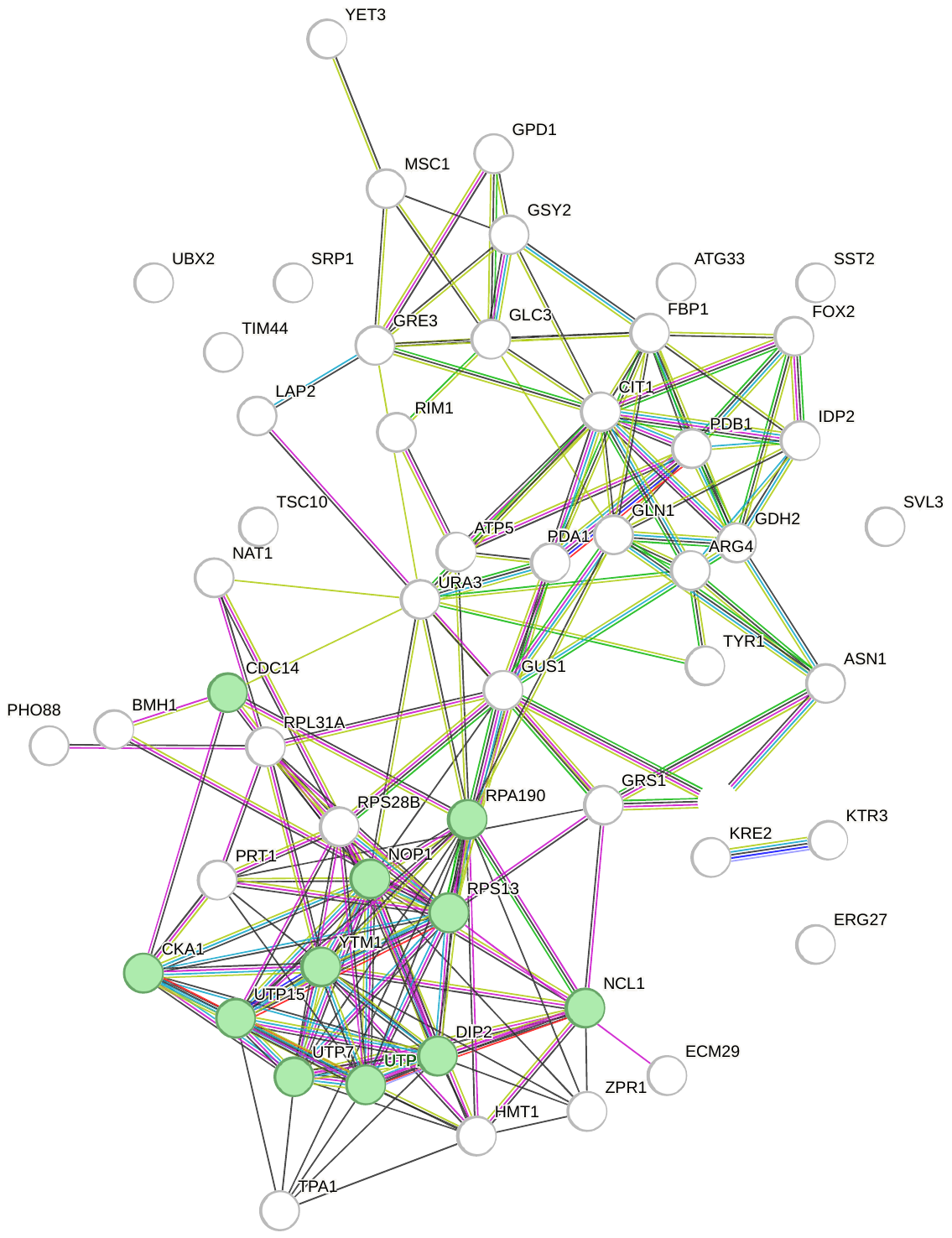**B.** |
| --- | --- |
| **C.**  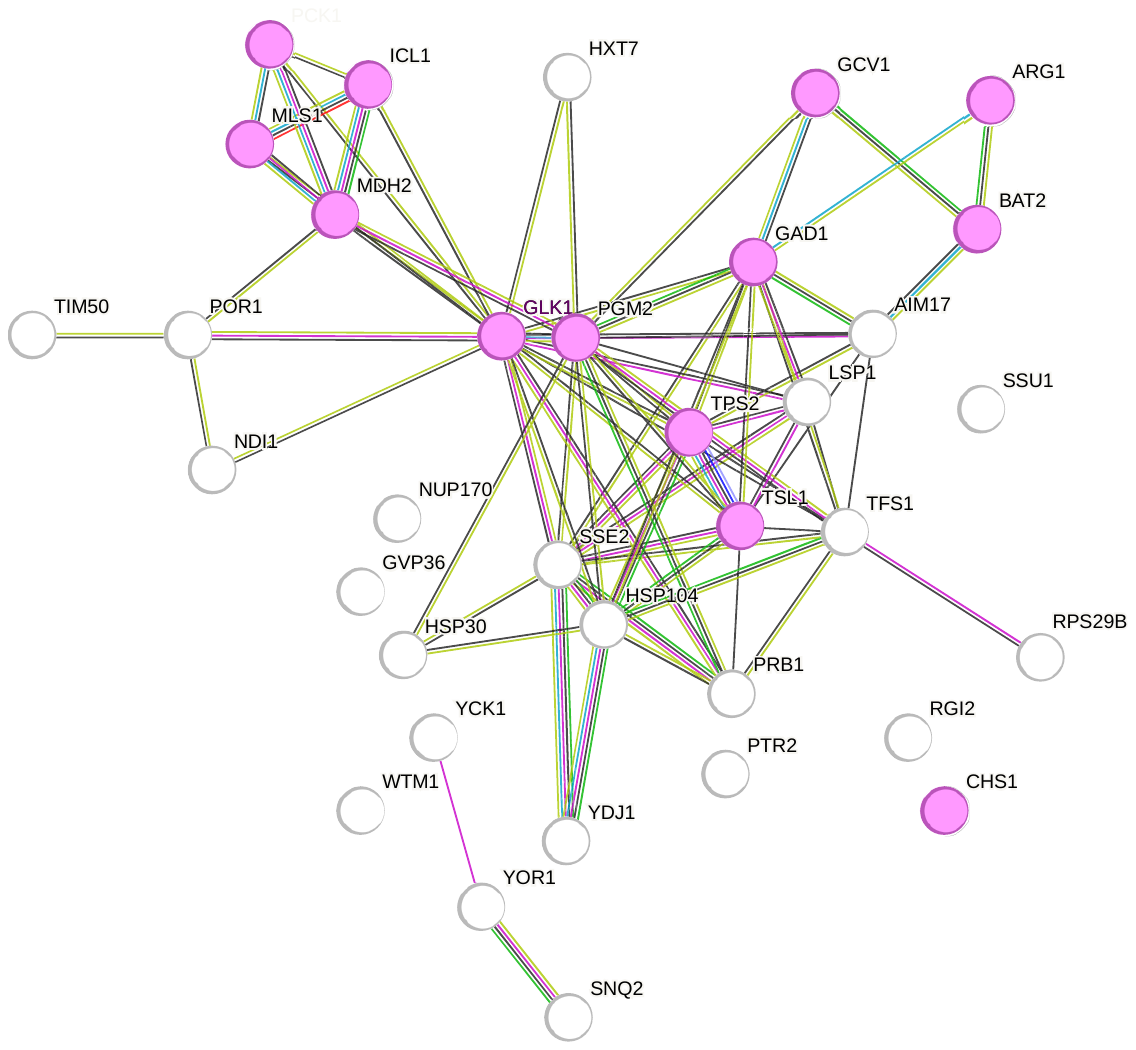 | |
| **SI Figure 4. Functional enrichment analysis of the target genes regulated by transcription factors specifically associated with either upregulated or downregulated proteins using STRING-DB. A.** Enrichment network of target proteins that are increasingly regulated by the specific transcription factors. Gene Ontology analysis of cellular compartments shows the enrichment of cell periphery related proteins (red nodes, 30 proteins, FDR = 0.02) and mitochondrial proteins (blue nodes, 45 proteins, FDR = 8.9e-5) **B.** Enrichment network of target proteins that are decreased by the specific transcription factors. Gene Ontology analysis of cellular compartments shows the enrichment of nucleolus related proteins (green nodes, 11 proteins, FDR = 0.02) **C.** Enrichment network illustrating proteins that are commonly regulated by specific transcription factor in both upregulation and downregulation. Proteins highlighted in magenta represent the cluster involved in metabolic pathways (13 proteins, FDR = 0.002). | |

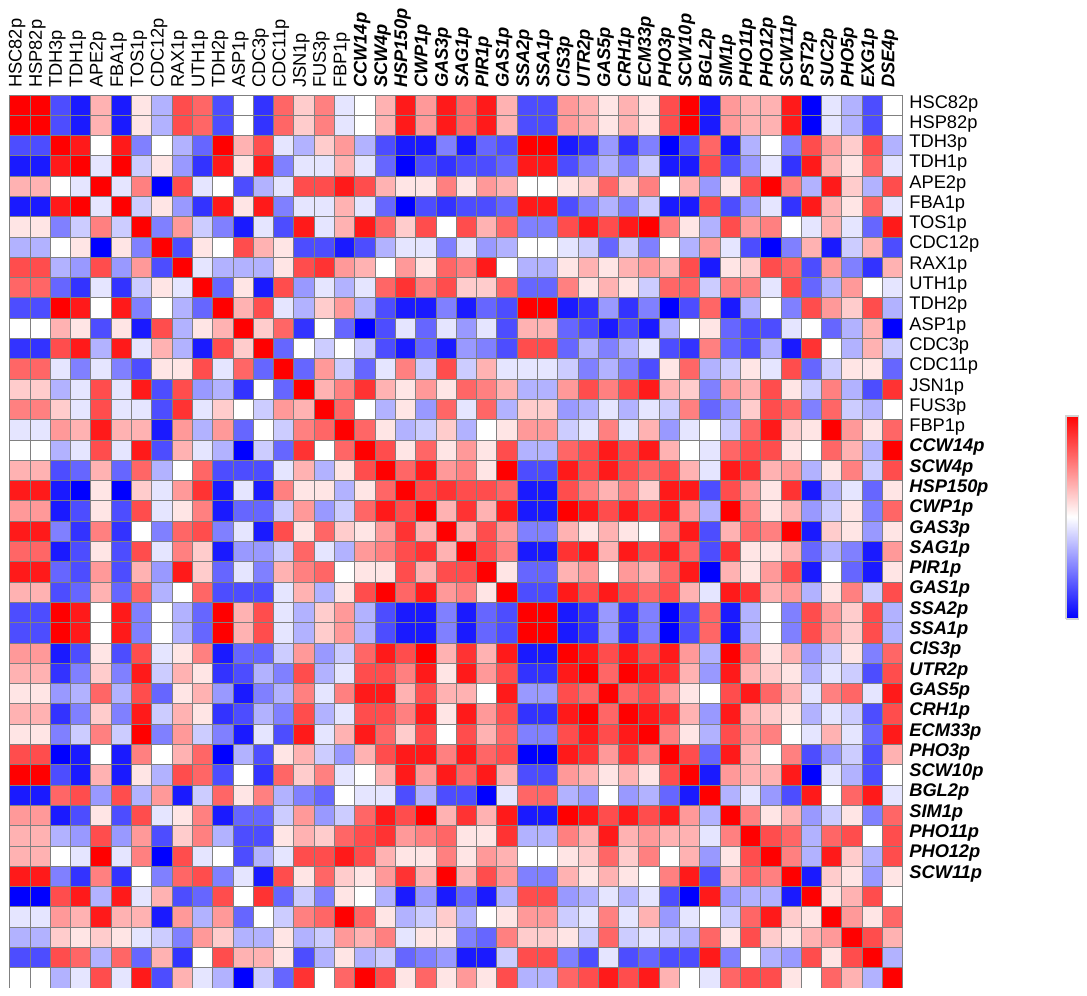
**A.**

**B.**

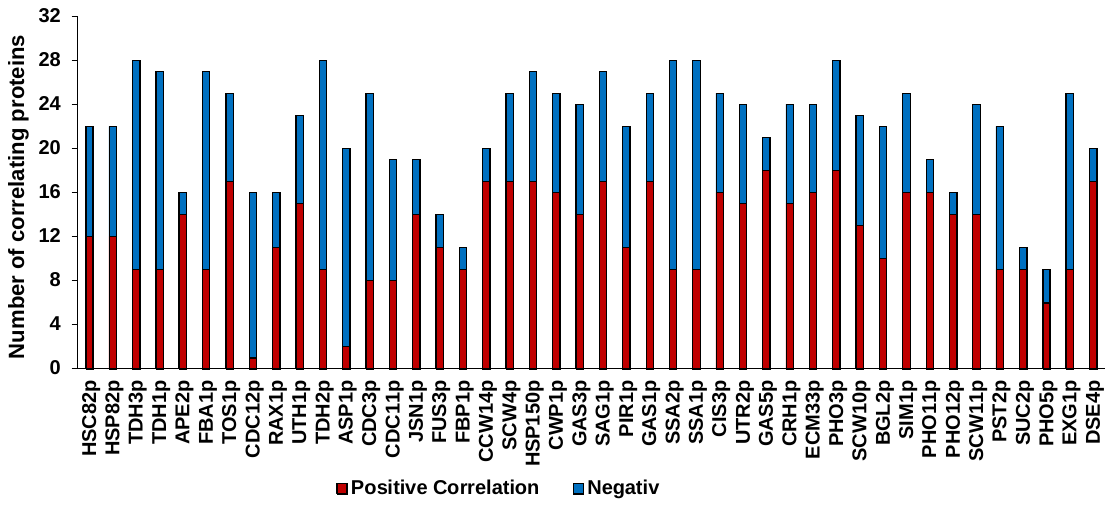

**SI Figure 5. Spearman rank correlation of CWPs expression during the kinetics of batch culture of S288c strain in bioreactor from our dataset, using Morpheus software. A.** Correlation matrix of different CWPs expression during the batch culture. The scale for correlation coefficient ranges between – 1 (dark blue) and 1 (dark red), centered around 0 (white). **B.** Statistics of positively correlated (red bars, correlation coefficient > 0.5) and negatively correlated (blue bars, correlation coefficient < -0.5) per CWP.

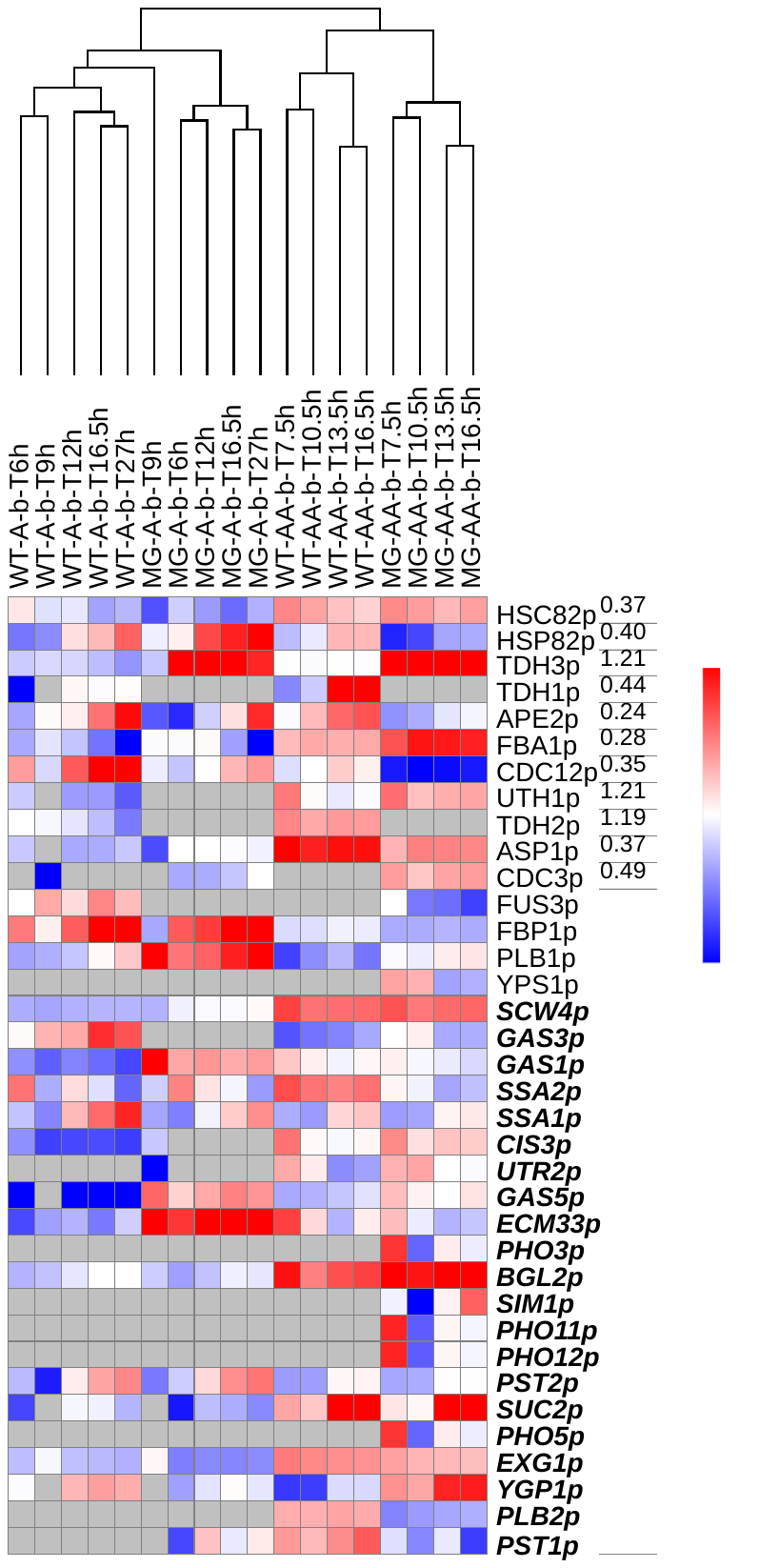

**SI Figure 6. Hierarchical clustering heatmap by Euclidean distance for Z-score normalized protein abundances established with Morpheus software, showing time point-specific expression signatures of CWPs in *S. cerevisiae* grown in batch mode with a rich medium in bioreactor from den Ridder *et al* dataset. (22)** The dataset is acquired for CEN.PK113-7D wild-type strain (WT) and minimal glycolysis mutant (MG), cultured in aerobic (A) for 30 h or in anaerobic (AA) for 24 h in label-based bottom-up quantitation (TMT 10-plex). The scale for z-score ranged between -2 (dark blue) and 2 (dark red), centered around 0 (white). The grey color indicates that the protein was not identified / quantified in the sample.

**A.**

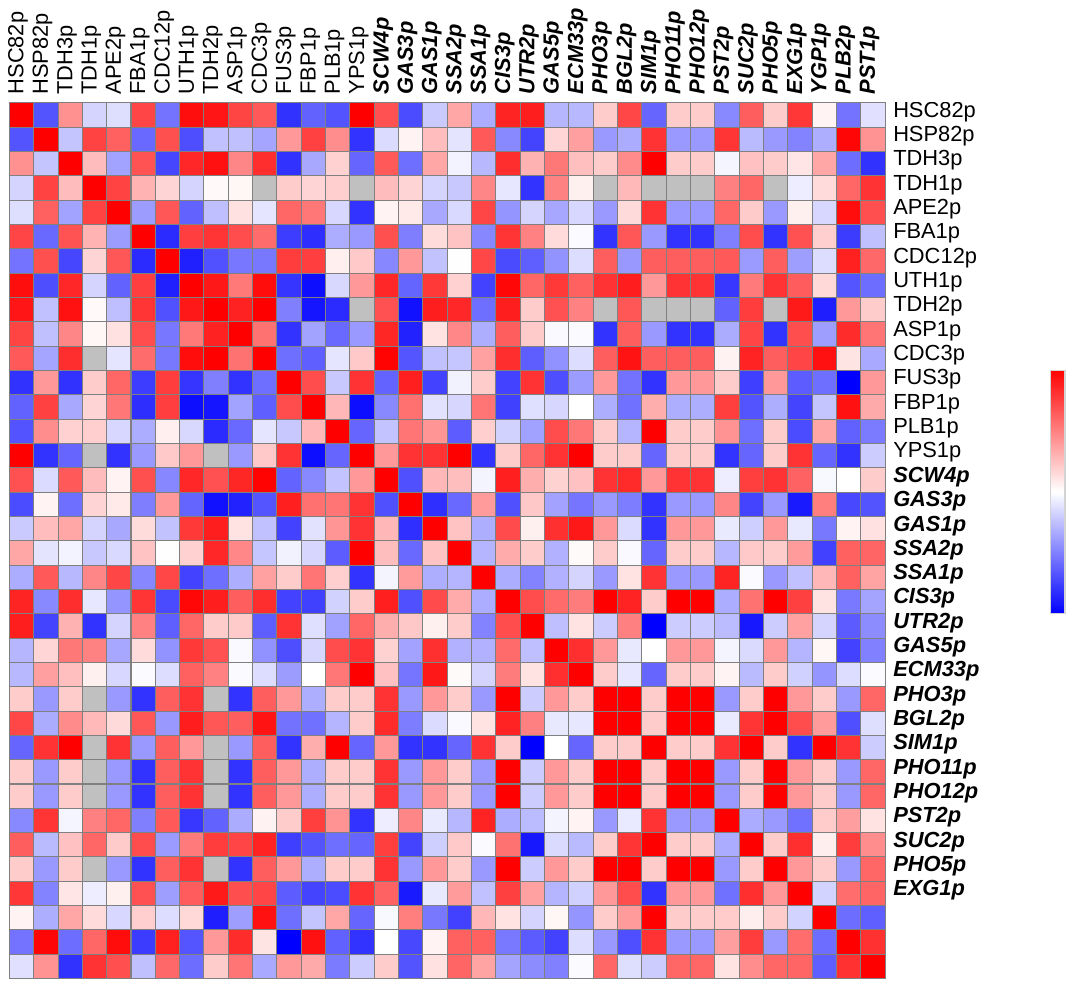

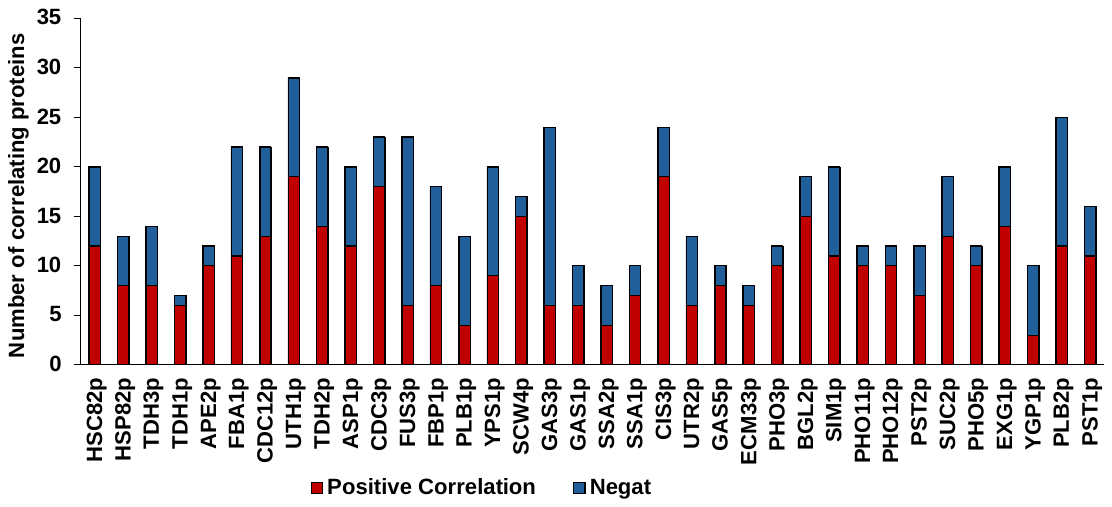
**B.**

**SI Figure 7. Spearman rank correlation of CWPs expression during the kinetics of batch culture of** **CEN.PK113-7D wild-type and mutant strains in bioreactor from den Ridder *et al* study, using Morpheus software. A.** Correlation matrix of different CWPs expression during the batch culture. The scale for correlation coefficient ranges between – 1 (dark blue) and 1 (dark red), centered around 0 (white). **B.** Statistics of positively correlated (red bars, correlation coefficient > 0.5) and negatively correlated (blue bars, correlation coefficient < -0.5) per CWP.

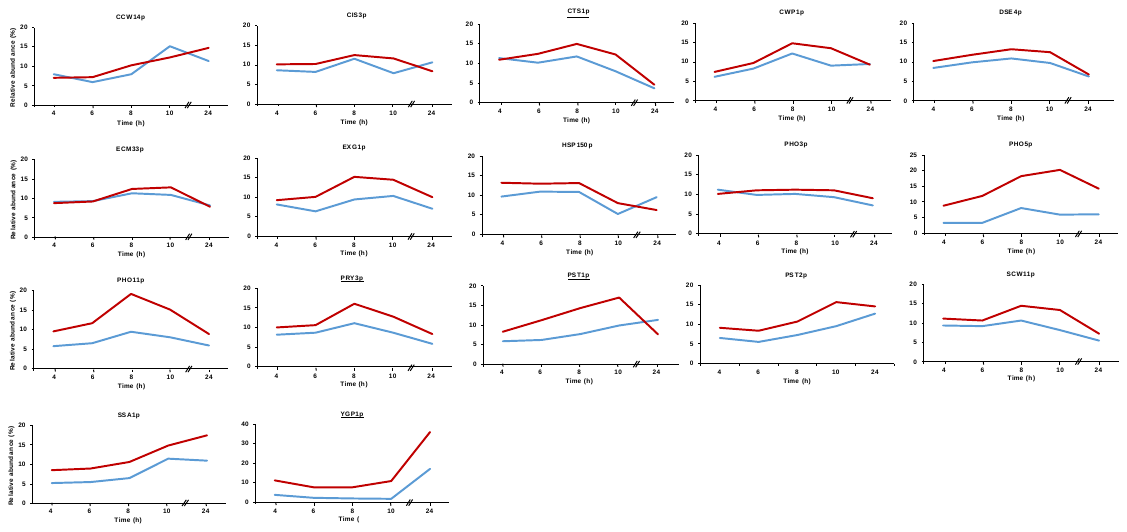
**A.**

**SI Figure 8. Relative abundance changes of strictly cell wall - localized CWPs during batch culture of the wild-type *S. cerevisiae* BY4742 strain derived from S288c strain and its Hap2p-deletion mutant in 1 L flasks. A.** CWP with Hap2-dependent expression. **B.** CWP with Hap2-independent expression. Data retrieved from Murphy *et al.* (21) represent mean relative abundances (%) from triplicate experiments. The blue curve shows protein expression in the wild-type strain, and the red curve shows expression in the Hap2p-deletion mutant. Underlined CWPs were quantified only in the Murphy *et al.* study (21).

**B.**

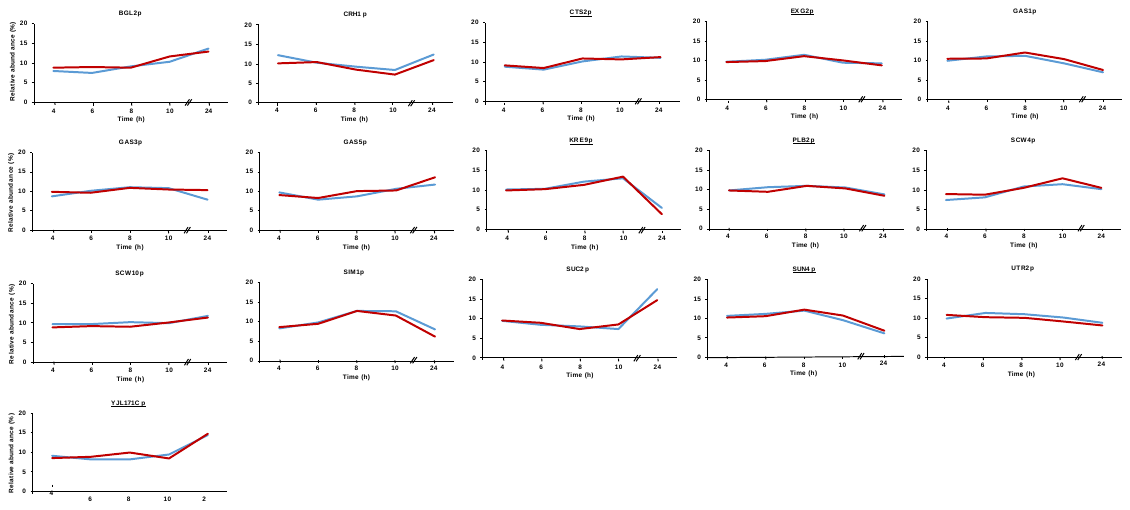

**SI Figure 8. Relative abundance changes of strictly cell wall - localized CWPs during batch culture of the wild-type *S. cerevisiae* BY4742 strain derived from S288c strain and its Hap2p-deletion mutant in 1 L flasks. A.** CWP with Hap2-dependent expression. **B.** CWP with Hap2-independent expression. Data retrieved from Murphy *et al.* (21) represent mean relative abundances (%) from triplicate experiments. The blue curve shows protein expression in the wild-type strain, and the red curve shows expression in the Hap2p-deletion mutant. Underlined CWPs were quantified only in the Murphy *et al.* study (21).

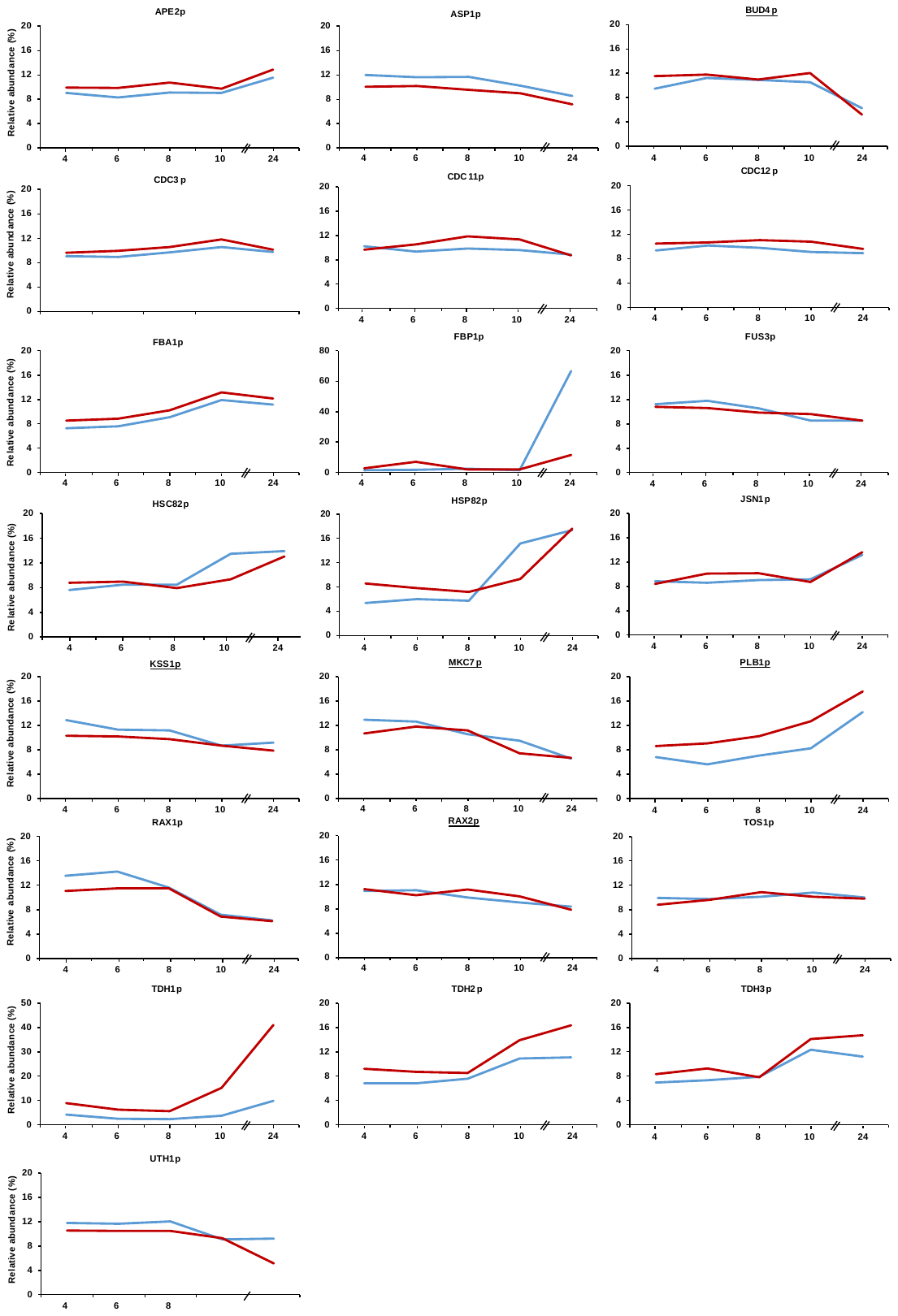

**SI Figure 9. Relative abundance changes of CWPs that are also localized in other organelles during batch culture of the wild-type *S. cerevisiae* BY4742 strain derived from S288c and its Hap2p-deletion mutant in 1 L flasks from Murphy *et al*. (21).** Data retrieved from Murphy *et al.* (21) represent mean relative abundances (%) from triplicate experiments. The blue curve shows protein expression in the wild-type strain, and the red curve shows expression in the Hap2p-deletion mutant. Underlined CWPs were quantified only in the Murphy *et al.* study (21).

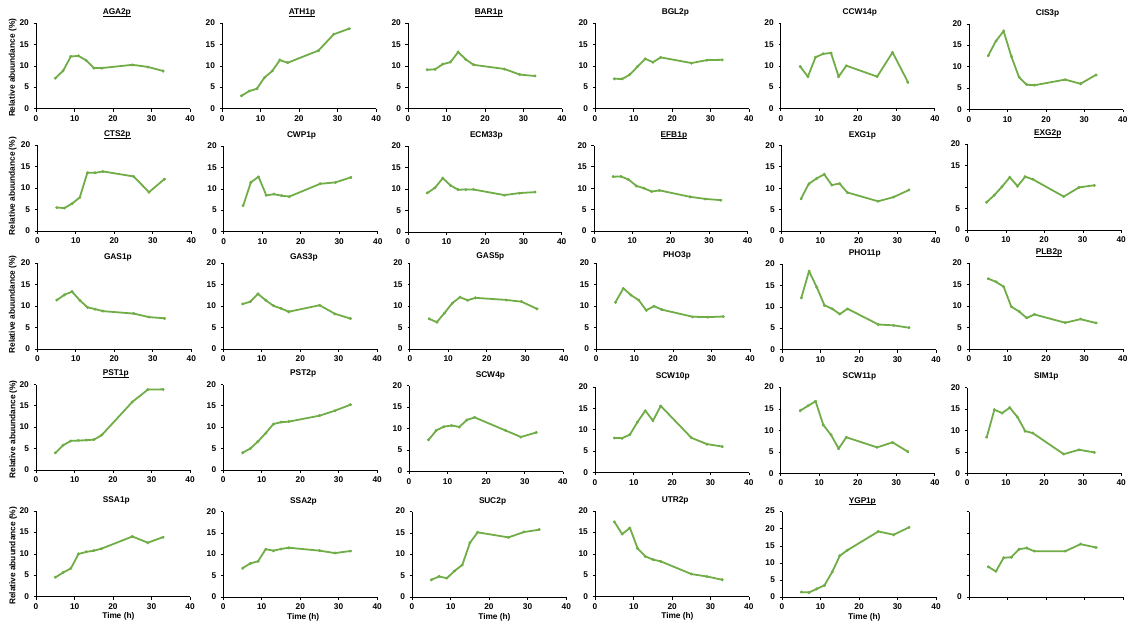

**SI Figure 10. Relative abundance changes of strictly cell wall – localized CWPs during batch culture of the wild-type *S. cerevisiae* DBY7286 strain in 1 L flasks for 33 h.** Data retrieved from Murphy *et al.* (21) represent mean relative abundances (%) from triplicate experiments. Underlined CWPs were quantified only in the Murphy *et al.* study (21)

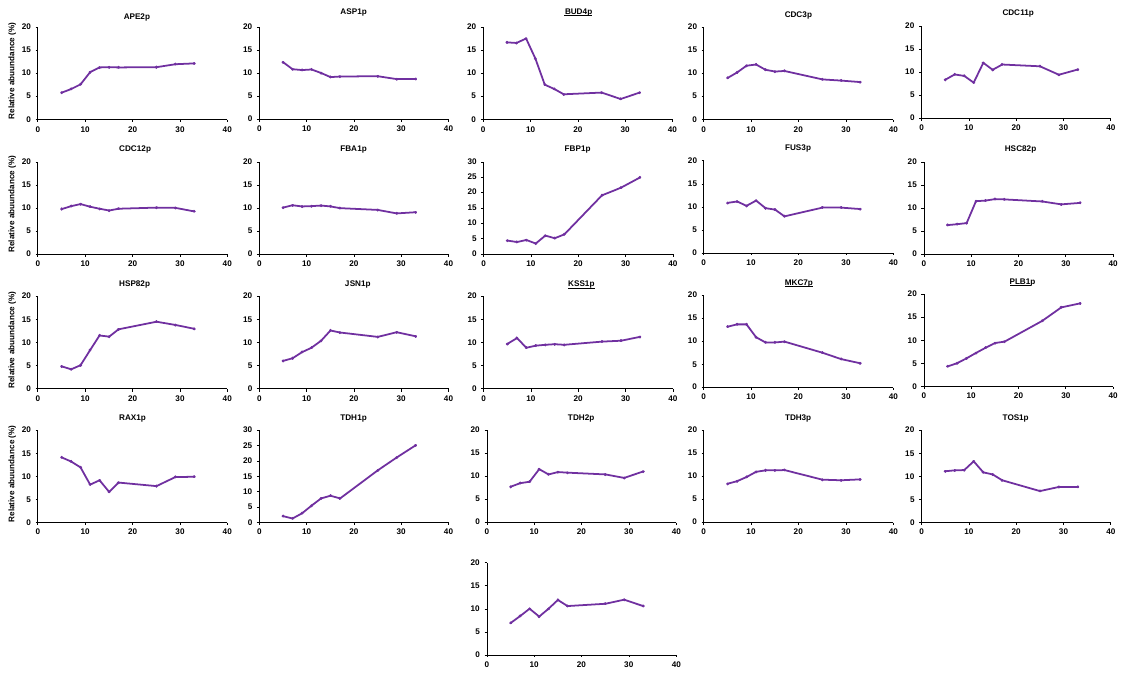

**SI Figure 11. Relative abundance changes of CWPs that can be in other organelles during batch culture of the wild-type *S. cerevisiae* DBY7286 strain in 1 L flasks for 33 h.** Data retrieved from Murphy *et al.* (21) represent mean relative abundances (%) from triplicate experiments. Underlined CWPs were quantified only in the Murphy *et al.* study (21).

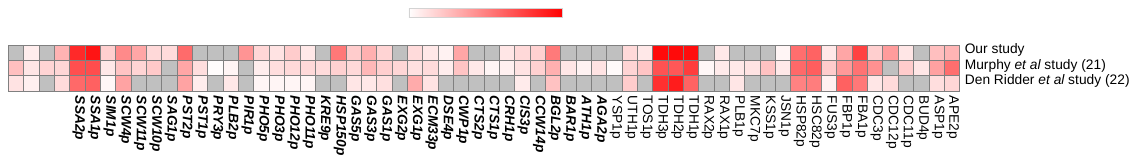

**SI Figure 12. Comparison of CWP sequence coverage (%) between our dataset and those reported by Murphy *et al.* (21) and den Ridder *et al.* (22).** For each study, the maximum sequence coverage value observed for each CWP across all experimental conditions is shown. CWPs exclusively localized to the cell wall are indicated in ***bold italics*,** while CWPs that may also occur in other organelles are shown in regular font. The grey color indicates that the protein was not identified / quantified in the study.
